# Structural conservation and divergence across the Receptor Tyrosine Kinase superfamily

**DOI:** 10.1101/2024.12.31.630944

**Authors:** Anna Fassler Bakhman, Rachel Kolodny, Mickey Kosloff

## Abstract

Members of the Receptor Tyrosine Kinase (RTK) superfamily are regulators of cellular signaling, playing essential roles in cellular growth, differentiation, and survival. Dysregulation of RTKs leads to diseases such as cancer, diabetes, and inflammatory disorders, making them important therapeutic targets. Despite extensive research on RTKs, the structural diversity and evolutionary relationships across the superfamily are not fully understood. Here, we systematically compared structural conservation and divergence among 245 extracellular domains from 54 RTKs, across 18 RTK families. Using experimentally-resolved structures and AlphaFold2 models, we conducted an all-versus-all structural alignment to explore domain architecture and quantify significant structural similarities within the RTK superfamily. We curated a comprehensive database encompassing PDB structures, 3D folds, ligand-binding properties, and sequence information of all RTK domains analyzed (https://fasslero.github.io/RTK-domains). Our analysis revealed numerous inter-family similarities and remote evolutionary connections, in particular among ligand-binding domains (LBDs), and distinct structural domain types and unexpected dissimilarities among domains previously classified as related. Our work highlights the intricate balance between structural conservation and divergence in RTKs and sheds light on the evolutionary mechanisms shaping this critical superfamily.

## Introduction

The receptor tyrosine kinase (RTK) superfamily comprises a diverse group of cell-surface receptors essential for normal development and processes like cellular growth, differentiation, and homeostasis (1-4). Dysregulated RTK activation can result in various pathologies, including cancer, diabetes, inflammation, and tumor angiogenesis – making RTKs central therapeutic targets (4-7). RTKs have been classified into 20 families, primarily based on sequence similarity and structural characteristics of the extracellular regions (4, 8) }(2, 3). 18 out of these 20 RTK families feature an extracellular region, which mediates ligand binding and/or activation: VEGFR, PTK7, PDFR, FGFR, Tie, Axl, Trk, MuSK, ROR, MET, ROS, Ryk, RET, InsR, EphR, DDR, ALK, and ErbB. while all 20 RTKs contain a single transmembrane helix and a cytoplasmic region that includes a conserved tyrosine kinase domain, the two families that lack extracellular regions and consist only of transmembrane and intracellular domains are LMR and STYK1 (2, 9). The extracellular region of RTKs can vary in length and in structural elements and plays a key role in ligand-induced dimerization, that, in turn, activates intracellular signaling. Activation leads to autophosphorylation and initiates downstream signaling pathways via protein recruitment and protein phosphorylation (1, 9-12). While the intracellular kinase region is conserved across the superfamily (13), the extracellular domains exhibit greater diversity, reflecting adaptation to dissimilar ligands. Within each RTK family, the number of extracellular domains and their 3D folds are often conserved among members, with the exception of the two members of the ALK family, which have six and two extracellular domains (14).

Lemmon *et al*. classified the extracellular regions of RTKs according to 19 distinct structural domains that are found among these 18 RTK families (2). Among these 19 distinct domains, the most common structural domains were immunoglobulin-like (Ig-like) domains, appearing in ten of the 18 RTK families. Ig-like domains are characterized by a two-layered β-sheet structure in a “sandwich” configuration and often function as ligand-binding domains (LBDs) – playing crucial roles in receptor-ligand recognition (15). Fibronectin type III (FN3) domains, which are present in four RTK families, have a similar 3D fold to Ig-like domains and their β-sheets can be superimposed (16). On the other hand, 13 out of the 18 RTK families contain 17 additional structural domains that were classified as distinct from one another and dissimilar to Ig-like and FN3 domains (2). Among the ten RTK families that contain Ig-like domains, the extracellular regions of the VEGFR, PTK7, PDGFR, and FGFR families are composed exclusively of Ig-like domains, while six other families feature a combination of Ig-like and other domain types. Furthermore, previous studies suggested that the structurally-similar PDGFR and VEGFR, as well as FGFR, share a common ancestor (2, 17). Indeed, numerous cross-family RTK-ligand interactions have been identified between PDGFR ligands and VEGFR receptors, as well as between VEGF ligands and PDGFR receptors (18). Based on these results, Mamer *et al*. suggested that such ligand-receptor cross-talk enables functional overlap across the entire RTK superfamily. Notably, three RTK families bind ligands via extracellular domains that are found only in a single RTK family (2). These include the discoidin domain in the DDR family, the TNFL-GR domain in ALK, and the jelly-roll domain in EphR (14, 19, 20). Each of these domains has distinct 3D structural characteristics – the discoidin domain forms a β-barrel, the LTK LBD features a TNF-like β-sandwich, and the Eph LBD adopts a jellyroll β-sandwich, illustrating the structural diversity across RTK families (14, 21-23). Differences in RTK extracellular structural components were also suggested to be key contributors to ligand-binding specificity (24, 25). Yet, despite extensive research, questions remain about how structural variations across RTK families are linked to functional roles and how evolutionary processes have shaped these relationships. While previous studies have provided insights into the structure and activation mechanisms of individual RTKs, a comprehensive understanding of the evolutionary connections between structure and function remains elusive.

In this study, we investigated the structural relationships across the RTK superfamily by comparing all 245 extracellular structural domains in the 18 RTK families that contain such regions. Using both experimental structures and AlphaFold2 models, we conducted an all-versus-all structural alignment, assuming they are spread across a continuum of protein structure space (26). We search for inter-family similarities, particularly across distinct ligand-binding domains from different structural domain types, and for potential new evolutionary relationships within the RTK superfamily. To complement this, we integrated functional data on ligand-binding domains, focusing on receptors with known ligands or established ligand-binding domains. We curated a database encompassing all 245 domains from 54 individual RTKs analyzed here, accessible at: https://fasslero.github.io/RTK-domains. This database provides all 3D domain structures analyzed, 3D fold classifications for each domain, ligand-binding domain categorizations, and all sequence data. This resource and our results lay the groundwork for future studies aimed at unraveling the complexities of RTK-mediated signaling and evolution and better targeting RTKs with more specific therapeutics.

## Results

To quantify structural similarities across the RTK superfamily, we analyzed the extracellular region of each receptor we identified in UniProt (see Materials and Methods). We classified these regions based on the literature, separating them into distinct structural domains based on domain boundaries from the ECOD database (27, 28), totaling 19 structural domain types (Fig. 1). We note that four of these domains were classified differently in previous studies (2) – the IPT and jelly-roll domains were previously classified as Ig-like and Ephrin-binding, respectively, while the heparin-binding and TNFL-GR domains were both previously classified as Mam domains. In addition, we classified two domains in the ErbB family as EGF domains based on their classification in ECOD, while previously they were classified as cysteine-rich domains (2). We re-classified these domains based on our visual inspection, ECOD data, and our domain boundaries analysis, revealing key differences from previous classification. Papers that showed new structural data further supported this re-classification (20, 29). We separated the extracellular region of EphR into four domains that include two FN3 domains based on (20), instead of the classification of two distinct FN3 domains and one ephrin-binding domain in (2). We also separated the two members of the ALK family, ALK and LTK, into six and two domains, respectively, based on (14), rather than three distinct domains as in (2). Our resulting dataset encompasses 54 receptors, divided into 245 distinct structural domains across these 18 RTK families. Each domain was labeled according to the ECOD database and/or the literature (Fig. 1, Supplementary Table 1). The ECOD domain classification conflicted with the literature in the RET, InsR, EphR, DDR, and ALK families – in these cases we prioritized the literature classification. 52% of these domains (128 out of 245) did not have an experimentally determined structure in the Protein Data Bank (PDB), and thus we used AlphaFold2 (30) to predict their structures (Supplementary Fig. 1). In five RTK families, AlphaFold2 was used to predict the entire structure of all family members, while in ten RTK families we predicted only the specific domains that were missing in the structures. The five families lacking experimental structures in the PDB were PTK7, ROR, MET, ROS, and Ryk. For these RTKs, domain boundaries were defined using a combination of literature, AlphaFold2 Predicted Aligned Error (PAE) output, and visual inspection of the predicted models – similar to the approach described in (31). The goal of this integrated approach was to accurately and consistently identify domain boundaries, even in the absence of experimental data.

**Figure 1.**
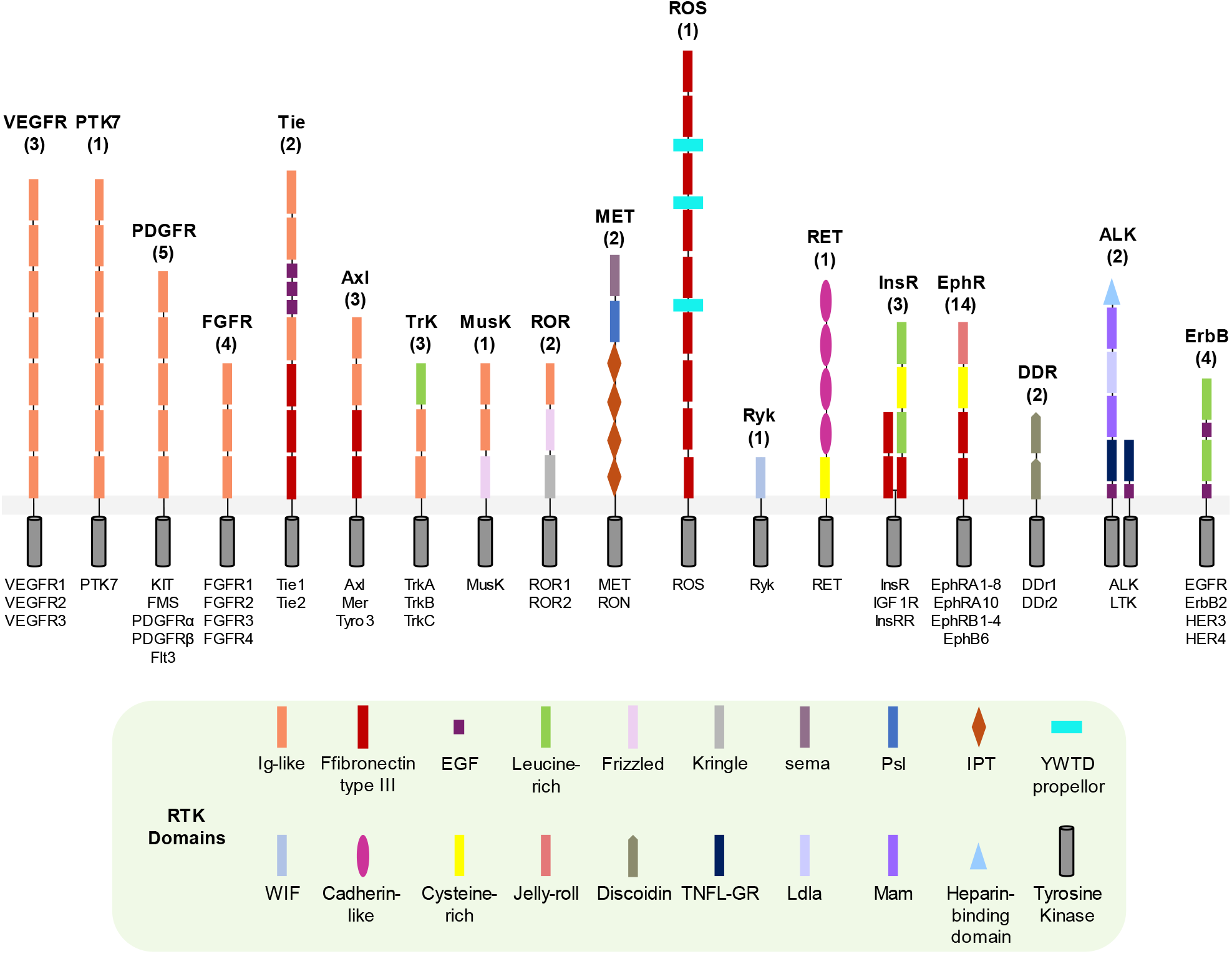
Classification of RTK extracellular domains across 18 receptor tyrosine kinase families. 54 human RTKs have been classified into 18 families, with extracellular regions illustrated schematically according to structural domains based on the literature and the ECOD server. Family members are listed beneath each receptor, with the number of receptors of each family indicated in parentheses above the receptor. Structural domains are labelled according to the provided key, while intracellular regions are marked as grey cylinders. The cell membrane is depicted as a light grey bar. The ALK family shows two different receptor schemes, reflecting differences in the number and composition of structural domains between the two ALK family members – ALK and LTK.

Next, we used TM-align (32) to structurally align all pairs within the set of 245 RTK domains. Since TM-scores may differ when comparing two proteins, depending on which protein is marked “query” and which “subject”, we calculated both directions and selected the higher TM-score (max-TM-score) of the two. We rely on the thresholds outlined by (32), where a TM-score > 0.5 implies that the two proteins share a similar topology (33), and a TM-score ≥ 0.8 indicates that the proteins certainly have the same topology. Using Cytoscape (34) and Cytostruct (35), we mapped all substantial similarities of TM score ≥ 0.5 onto a network. These similarities were further divided into two subsets: 0.8 ≤ max-TM-score ≤ 1 and 0.5 ≤ max-TM-score < 0.8 (Fig. 2).

**Figure 2.**
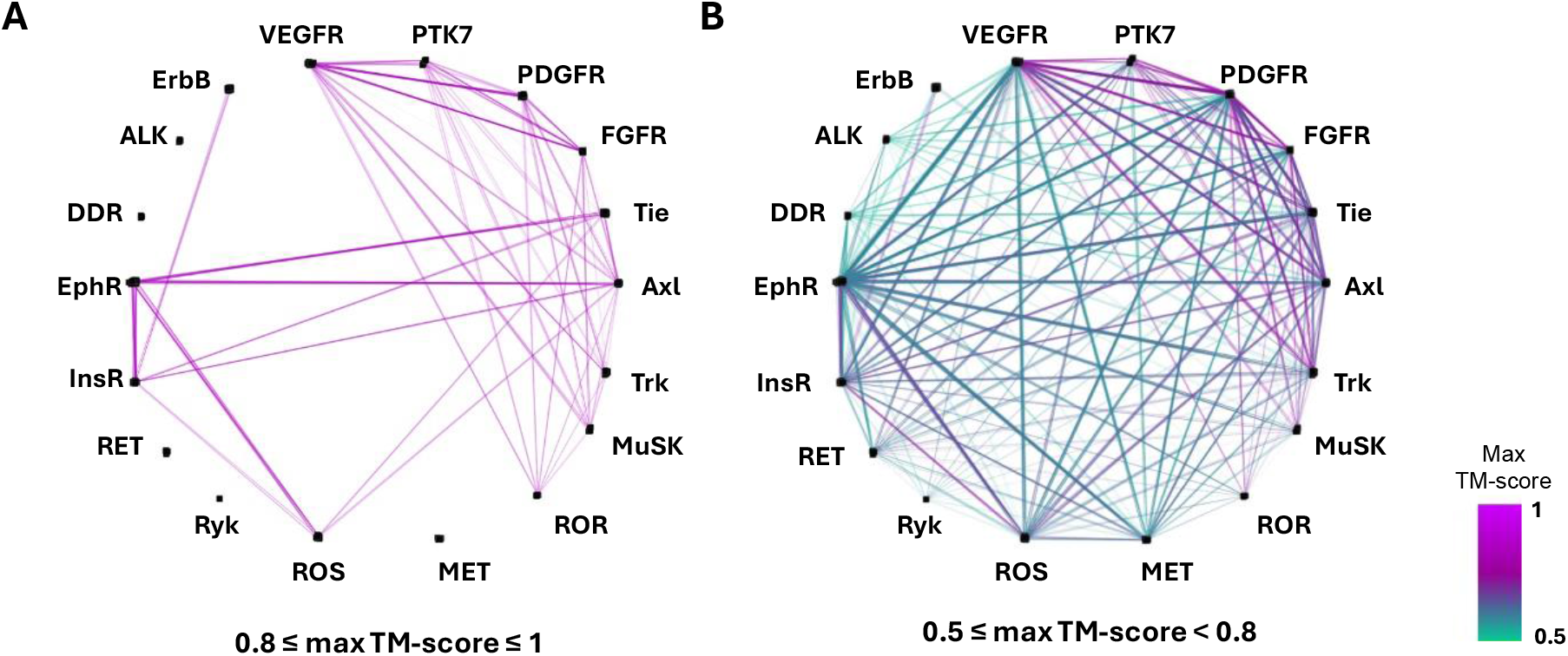
Structural similarity networks across 245 extracellular domains of 18 RTK families show wide-spread structural similarities between all families. **A**. Structural similarities among the 18 RTK families with 0.5 ≤ TM-score < 0.8. Each node represents an individual family, and edges indicate similarities between (at least one) members of the family and (at least one) member of other RTK families, with edge colors corresponding to the max-TM-score (see scale on right). **B**. Structural similarities among the 18 RTK families with 0.8 ≤ TM-score ≤ 1, illustrated as in panel A.

Within the higher TM-score range (0.8 ≤ max-TM-score ≤ 1), the number of interfamily similarities is lower than the number of structural similarities identified in the range of 0.5 ≤ max-TM-score < 0.8. At the higher structural similarity threshold, 13 of the families are sparsely connected through structure similarities to 1-11 other families (Fig. 2A), with ErbB showing the fewest and Axl showing the highest number of connections to other families. The remaining four families do not show structural similarities to other families at this higher range of TM-scores. As expected from previous studies, these high-similarity interfamily connections are dominated by families composed primarily of Ig-like and fibronectin type III (FN3)-like domains (Fig. 2A). On the other hand, the connection of the ErbB family to the InsR family, observed only between their leucine-rich domains, does not extend to other families.

Conversely, at the 0.5 ≤ max-TM-score < 0.8 range, all RTK families are densely connected to 3-17 other families (Fig. 2B). In fact, 12 out of the 18 families are connected through structure similarities to all other 17 families, while five families are connected to 13-16 other families. ErbB, which is connected to the fewest number of families, is connected to only three families – InsR, EphR, and Trk. Unexpectedly, most of these interfamily similarities are not only among families containing domains previously considered similar, Ig-like and FN3, or between domains classified as having the same 3D fold. Rather, most of these connections are between domains classified as having different 3D folds, and even across domains that were classified as distinct domains that appear exclusively only in one RTK family.

Within the unexpected interfamily similarities we identified, notable connections were observed between the ligand-binding domains (LBDs) of three structurally-distinct and dissimilar families – EphR, DDR, ALK – and between three additional LBDs – leucin-rich, FN3, and Ig-like (Fig. 3). Notably, two RTKs, PTK7 and ROS, are “orphan receptors” (25), for which no ligands have been identified, and for two RTKs, ROR and MuSK, the precise location where ligands bind is unknown. We also note that no ligand-binding characterization was performed for RTKs lacking a crystal structure of a ligand-receptor complex and for which data on ligand-binding domains was not available in the literature, which may lead to underestimation of similarities among LBDs. The Discoidin LBD of the DDR family, which was previously classified as structurally distinctive, is in fact structurally-similar to the Ig-like LBDs of the Axl, TIE, VEGFR, TRK, and FGFR families, as well as to the jelly-roll LBDs of the EphR family and the FN3 LBDs of the InsR family. Similarly, the jelly-roll LBD of the EphR family, another domain previously classified as structurally distinctive, is seen to be structurally similar to the Ig-like LBDs of the TIE, VEGFR, TRK, PDGFR, and FGFR families; it is also structurally similar to the Discoidin LBD of the DDR family and the FN3 domains of the InsR family. Additionally, the LBD of the ALK family, represented by the TNFL-GR domain that is not found in other RTKs, is structurally similar to the Ig-like LBDs of the TRK, PDGFR, Axl, VEGFR, and FGFR families.

**Figure 3.**
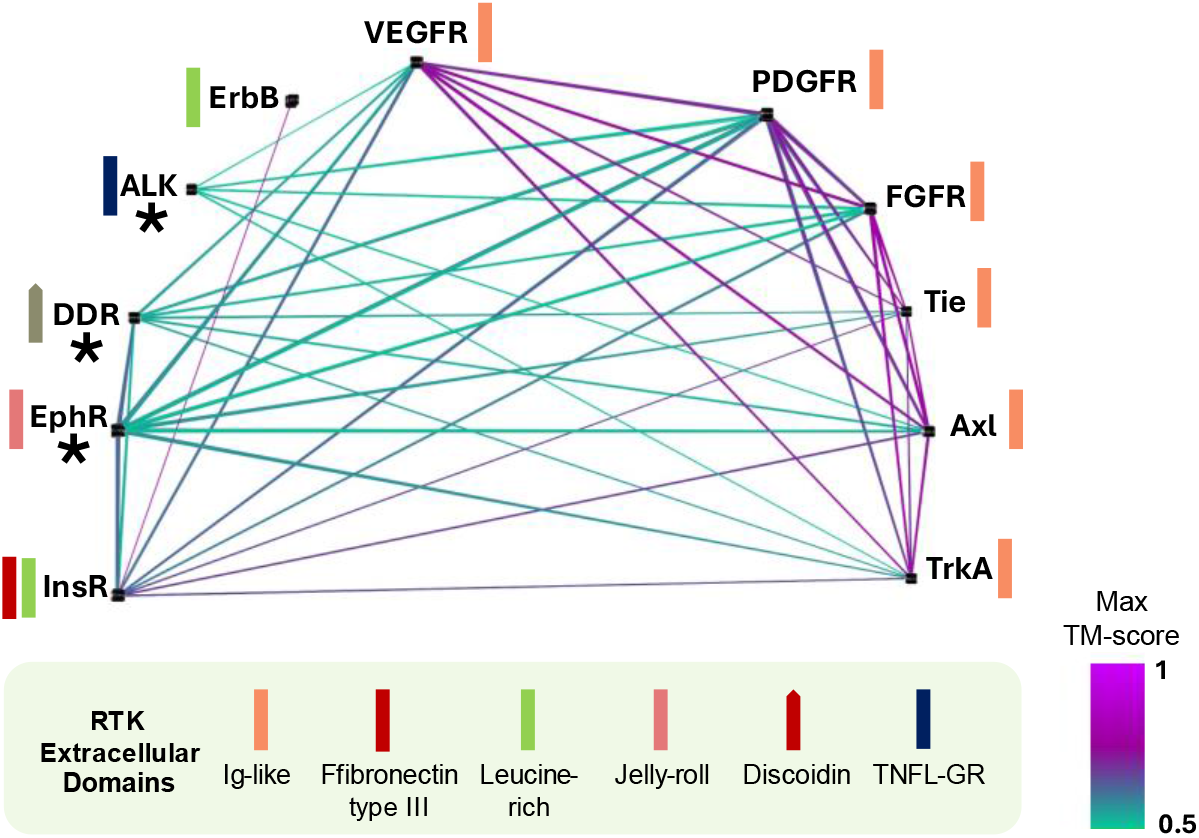
Ligand binding domains of 11 out of 18 RTK families are structurally similar to each other, including structural domains that are unique to individual families. Each node in the network represents an RTK family, and edges indicate structural similarities between (at least one) member of these families, with edge colors corresponding to TM-score values (see scale on right). The 3D fold of each ligand-binding domain in each RTK family is represented schematically, as in Fig. 1, next to the family name. Unique structural ligand-binding domains that are present exclusively in one RTK family are marked with a black asterisk.

To better identify patterns of structural conservation and divergence across the RTK superfamily, and to investigate intra-family similarities and differences, we mapped our results as a heatmap of per-domain similarity interaction (Fig. 4). As expected, intrafamily structural similarities are generally high across all RTK families (shown above the diagonal in Fig 4), with domains of the same type maintaining consistent structural conservation. In particular, VEGFR, PTK7, PDGFR, and FGFR, which are composed of exclusively Ig-like domains (Fig. 1), exhibit a high degree of intrafamily structural similarity across all domains, with TM-score values generally above 0.8. Similarly, the two DDR receptors containing Discoidin domains also display strong intrafamily structural conservation. Notably, in the ALK family, where ALK and LTK differ from one another in the number of extracellular domains (Fig. 1), intrafamily similarities are evident between these two dissimilar family members. The LBDs of these two RTKs are structurally identical with a TM-score of ∼1, despite the different domain composition.

**Figure 4.**
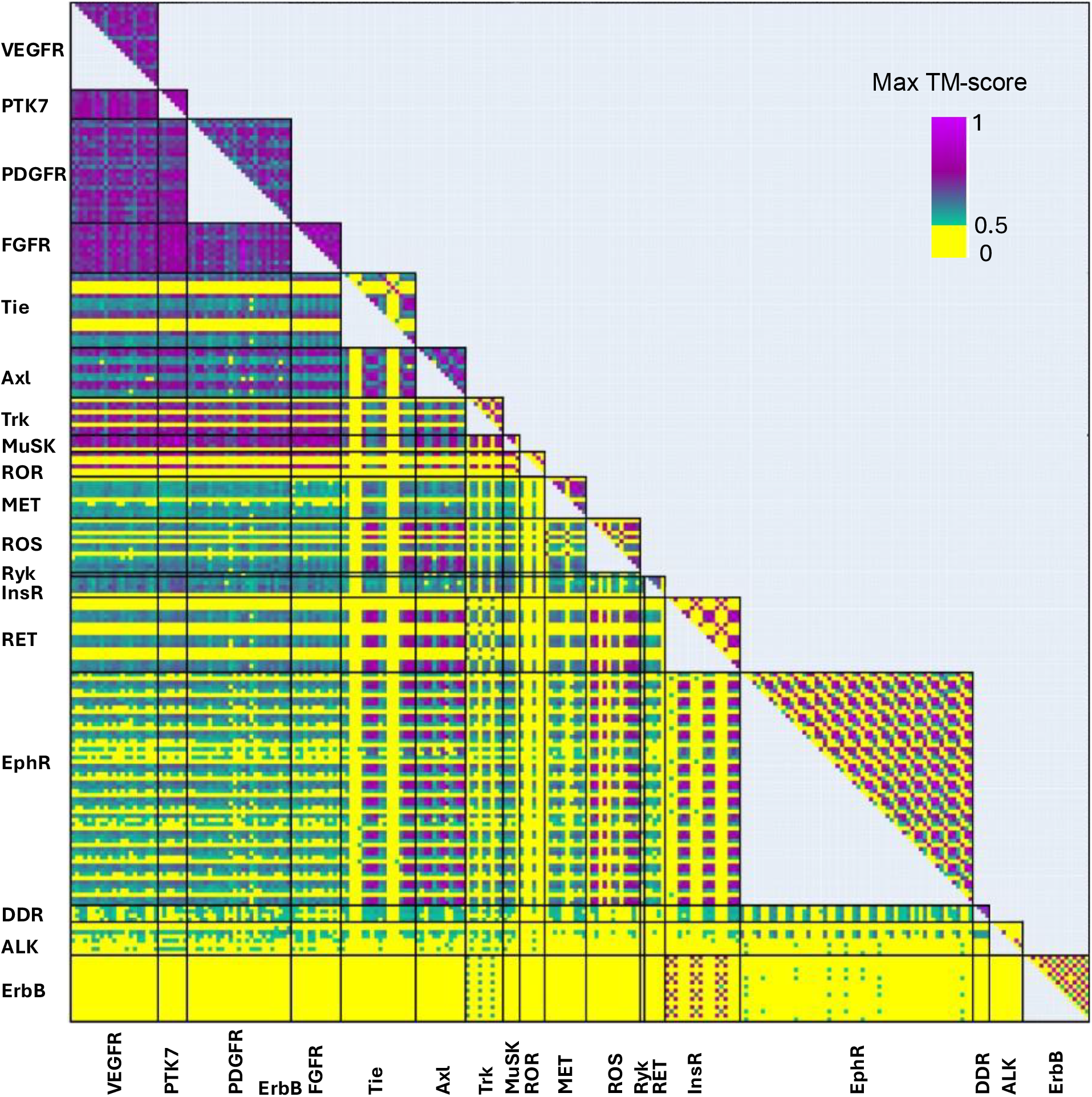
Pairwise intra- and inter-structural similarities across 245 structural domains in 18 RTK families, based on max-TM-score values. pairwise structural similarities among 245 domains. Rows and columns correspond to individual domains. Domains are grouped by the RTK family and ordered in an N’-to-C’ orientation. Intrafamily similarity interactions are shown above the diagonal, and interfamily similarity interactions are shown below the diagonal. The color gradient shows max-TM-score values, ranging from 0 to 1, where darker shades represent higher structural similarity and lighter shades indicate lower similarity (see scale at top right).

As shown above, substantial inter-family structural similarities are evident across the majority of the RTK superfamily (Fig. 4, under the diagonal). As expected, families composed exclusively of Ig-like domains, such as VEGFR, PTK7, PDGFR, and FGFR, exhibit substantial structural interfamily similarities – both among themselves and with other families that, in addition to other domains, include Ig-like domains: Tie, Axl, Trk, MuSK, and ROR. In addition to the structural similarity across Ig-like domains, families containing Ig-like domains also share substantial structural similarity with families containing FN3 domains, which were known to be structurally similar to Ig-like domains – Tie, Axl, ROS, InsR, and EphR. These interfamily similarities, particularly between Ig-like and FN3 domains, generally fall within a TM-score of ∼0.6-0.7 (colored light/dark-green in Fig. 4). Notably, these domains are typically LBDs (see above). Interestingly, both Ig-like and FN3 domains exhibit structural similarity with domains that were previously classified as distinct, including IPT, cadherin-like, discoidin, and Ryk domains, which are found in the MET, RET, DDR, and Ryk families, respectively. These structural similarities generally fall within a TM score of 0.5-0.6. Conversely, the LBDs in MET, RET, and Tie (Sema, cysteine-rich, and EGF domains, respectively), do not exhibit structural similarities to any other domains in the entire RTK superfamily (colored yellow in Fig. 4).

Among the leucin-rich and cysteine-rich domains, each group of domains substantial structural similarity. However, the exception is the cysteine-rich domain of the RET receptor that is not similar to any other cysteine-rich domain, nor to any other domain in the RTK superfamily. Regions of inter-family dissimilarity are also prominent between the ErbB / ALK families and all other RTK families. ErbB, which contains both EGF-like and leucine-rich domains, shows significant structural similarity only to the Trk, EphR, and InsR families. This limited connectivity stems from the leucine-rich domains shared among ErbB, Trk, and InsR, as well as the EGF-like domain in ErbB that is structurally-similar to the cysteine-rich domains of the EphR family. This global structural similarity is correlated with additional similarities, as both domain types are stabilized by intra-domain disulfide bonds and adopt compact globular folds that participate in similar functions of ligand binding and protein-protein interactions.

## Discussion

By combining experimentally-resolved structures with high-confidence AlphaFold2 predictions, we generated a comprehensive dataset of 245 domains spanning all 18 RTK families that have extracellular regions. We compared the geometries of all these extracellular domains by conducting all-versus-all structure alignment, identifying unexpected structural similarities and dissimilarities across the entire RTK superfamily. This analysis was integrated with data on ligand binding to gain insights into the structure-function relationships of the receptors. This was combined with a comprehensive and updated RTK classification of domain types based on extensive literature review, accessible at https://fasslero.github.io/RTK-domains.

Our results show that the RTK superfamily exhibits much denser and higher structural similarities, despite the diverse architectures reported for RTK extracellular regions. We observed expected similarities between RTK families with similar domain types as well as between families with domains that were previously classified as having distinctive 3D fold to those featuring Ig-like and FN3-like domains. Specifically, families that were classified as structurally distinctive, MET, Ryk, RET, EphR, DDR, and ALK. These families show structural similarity, both to families with Ig-like and FN3 domains, and to multiple other RTK families – highlighting a surprising degree of convergence among RTKs traditionally considered distinct. Moreover, structural similarities were identified among the LBDs of the DDR, EphR, and ALK families, specifically among the discoidin, jelly-roll, and TNFL-GR structural domains, respectively. These findings suggest that, despite their distinct domain architectures, these LBDs may share functional or evolutionary constraints that have driven convergent structural adaptations across RTKs. Indeed, “cross-talk” between the VEGFR and PDGFR families, where ligands from one family bind to receptors from the other, demonstrates the structural and functional flexibility within the RTK superfamily (18). Together with our broader results, these observations support a hypothesis of a shared evolutionary basis mediating diverse ligand-receptor interactions.

Our domain-level analysis revealed that divergence is evident among some RTKs with domains classified as architecturally distinctive. The ALK, RET, and ErbB families contain domains such as Mam and Ldla, WIF, cysteine-rich, and leucine-rich, which are found only in these families and exhibit very limited or no structural similarity to other RTK domains. For example, although RET, InsR and EphR contain domains that were previously classified as cysteine-rich domains, the cysteine-rich domain within the RET receptor is not structurally similar to the other RTK cysteine-rich domains found in the InsR and EphR families, or, in fact, to any domains across the RTK superfamily. Additionally, the ErbB family is structurally similar only to the Trk, InsR, and EphR families, reflecting their shared leucine-rich domains that in these families are LDBs. These divergences suggest evolution might have tailored these domains to the divergent ligands of each family.

Overall, our findings show a much higher degree of structural conservation across the RTK superfamily than previously anticipated. Some structural domains were used repeatedly by evolution for similar functions, such as Ig-like and FN3 domains, as recognized previously (16). However, the structural similarities observed among diverse ligand-binding domains, including the discoidin, jelly-roll, and TNFL-GR domains, suggest that these conserved architectures were shaped by evolution to optimize and maintain essential receptor-ligand interactions. These structural insights not only deepen our understanding of RTK evolution, but also provide a framework for investigating how structural diversity enables functional versatility. By leveraging our curated database and the comprehensive analysis presented here, future studies can systematically explore RTK structure-function relationships, offering critical insights into receptor biology and informing therapeutic innovations, with implications for understanding other receptor families as well.

## Acknowledgements

This work was supported by the United States-Israel Binational Science Foundation (BSF) and the United States National Science Foundation (NSF) grant number 2020627, by the International Development Research Centre (IDRC), the Israel Science Foundation (ISF), and the Azrieli Foundation grant number 3512/19, and by a grant from the Council for Higher Education through the Data Science Research Center at the University of Haifa.

## Materials and Methods

### Dataset collection

We searched UniProt (https://www.uniprot.org/) for all human sequences corresponding to the 18 RTK families identified as having extracellular structures, identifying 54 entries. These sequences were then used in BLAST searches at the NCBI website (https://blast.ncbi.nlm.nih.gov/Blast.cgi) to identify matching entries in the Protein Data Bank (PDB); 35 PDB IDs were thus identified as structures of extracellular regions of RTKs.

### Protein structures

Structures of RTKs with the following PDB IDs were included: KIT (2E9W), FMS (3EJJ). PDGFRa (7LBF), PDGFR (3MJG), Flt3 (3QS9), Axl (2C5D), Tyro3 (1RHF), TrkA (2IFG), Tie 2 (4K0V and 5MYA), InsR (6PXV), IGF1R (6JK8), InsRR (7TYM), MET (7MO7), RET (6Q2O), 7NX0 (ALK), 7NX4 (LTK), FGFR1 (1CVS), FGFR2c (2FDB), FGFR3 (3GRW), 7YSW (FGFR4), 5T89 (VEGFR1), VEGFR2 (2X1X and 3KVQ), VEGFR3 (4BSK), EphA2 (3FL7), EphA4 (4M4R), EphB2 (7S7K), EphB6 (7K7J), EGFR (1IVO), HER2 (1N8Z), HER3 (4LEO), HER4 (1N8Z), DDR1 (4AG4), DDR2 (2WUH), MuSK (2IEP and 3HKL). When multiple structures are available for a single receptor, we prioritized longer structures and those with better resolution.

### 3D structure prediction

We searched UniProt (https://www.uniprot.org/) to collect all human sequences of RTK proteins that lack an experimental structure. These sequences were used as queries in AlphaFold2 predictions using the ColabFold server (https://colab.research.google.com/github/sokrypton/ColabFold/blob/main/AlphaFold2.ipynb) (36).

### Structural alignment and network visualization of RTK domains

All-versus-all structure alignment was performed using TM-align (32). A custom script was developed to automate the process, systematically aligning all 245 RTK domains and extracting the TM-scores for further analysis. For each pair of aligned protein structures, the highest TM-score (max-TM-score) was selected for subsequent evaluation. All data from the all-versus-all structural alignment analysis was then imported into Cytoscape (v3.10.2) for network visualization (37).

