## Supplementary Figure 1 for "Structural conservation and divergence across the Receptor Tyrosine Kinase superfamily"

**VEGFR1**
**(VEGFR)**

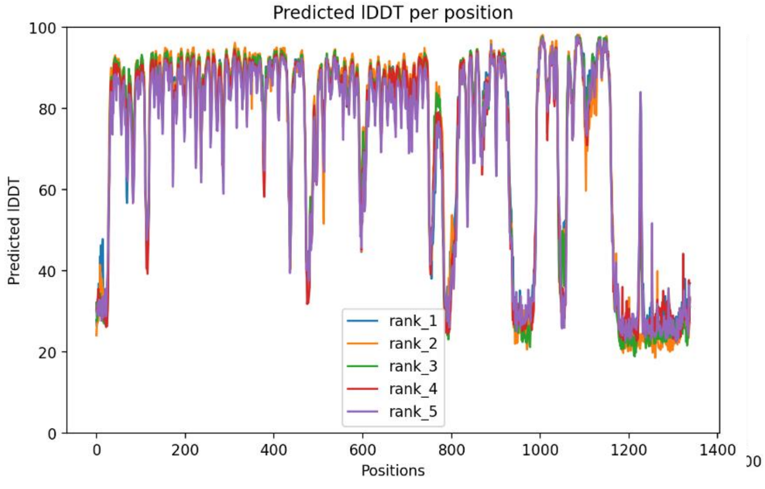


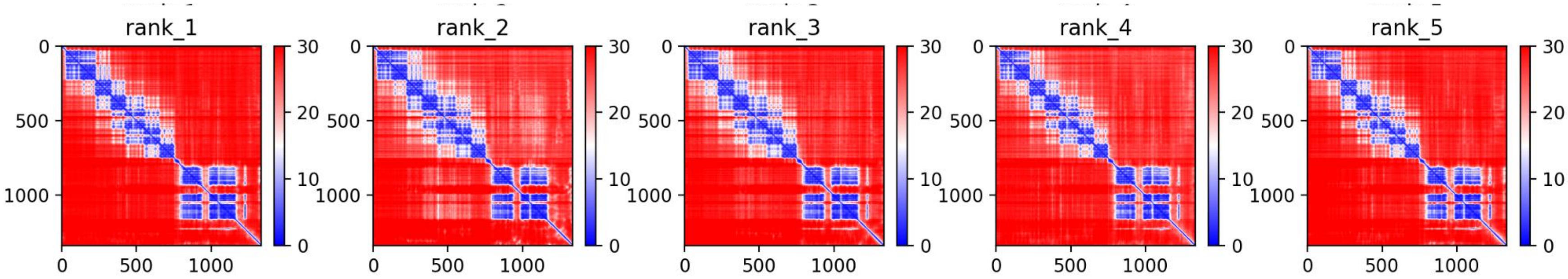

**VEGFR2**
**(VEGFR)**

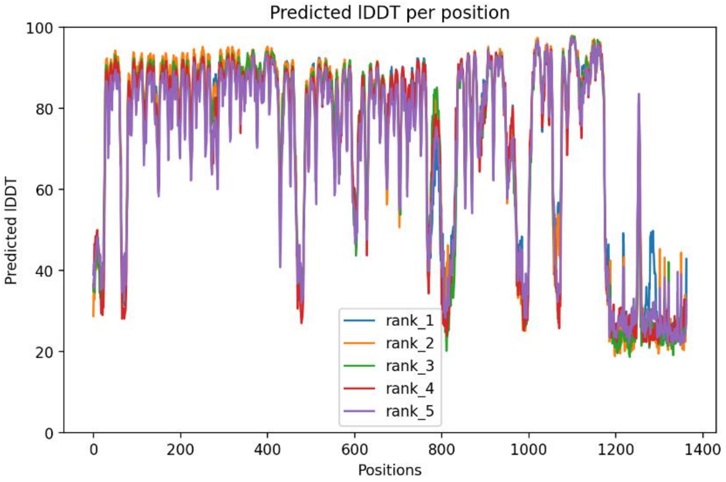


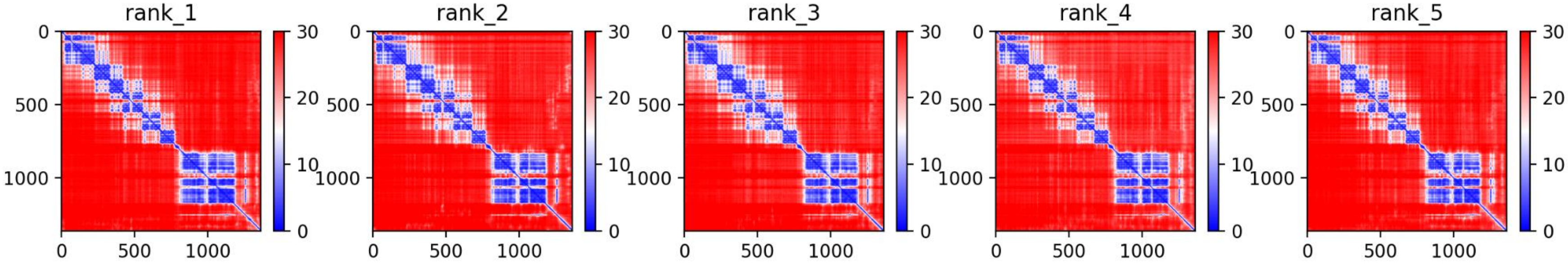

**VEGFR3**
**(VEGFR)**

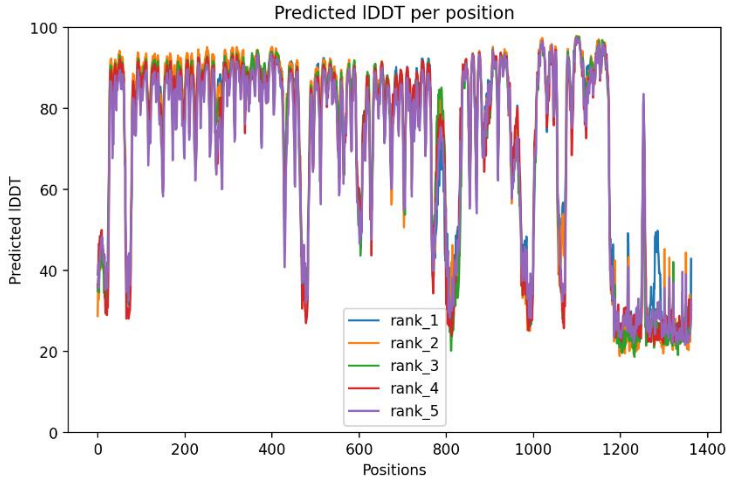


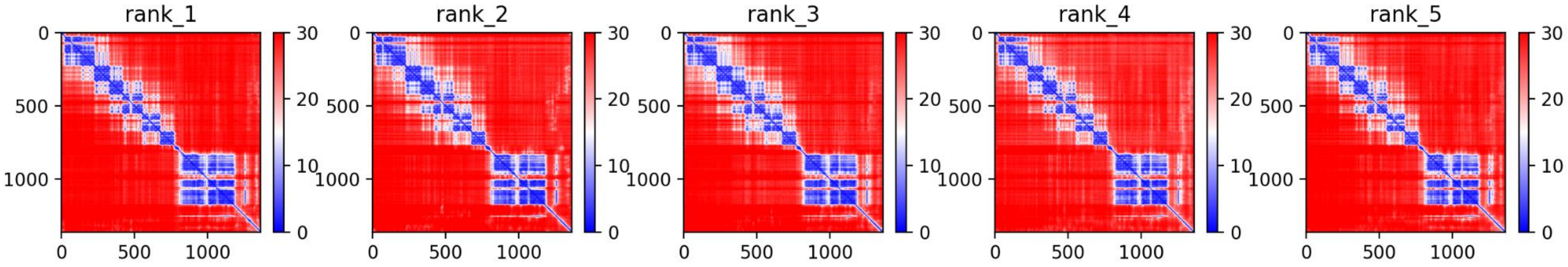

**PTK7**
**(PTK7)**

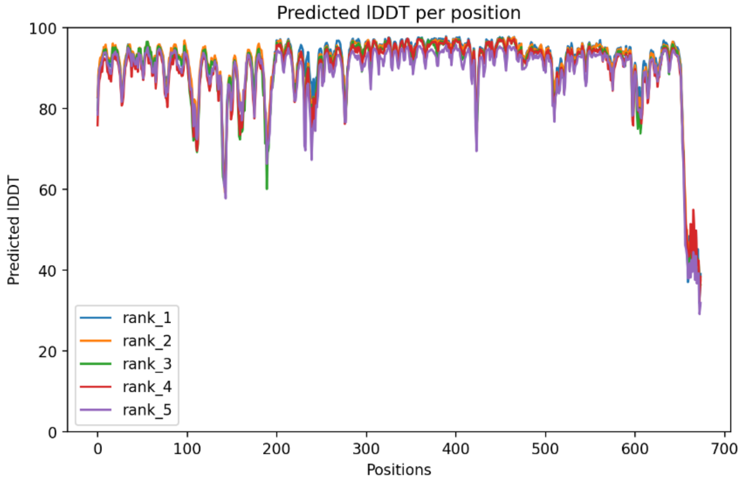

Predicted lDDT per position
Predicted lDDT
Positions
rank_1
rank_2
rank_3
rank_4
rank_5


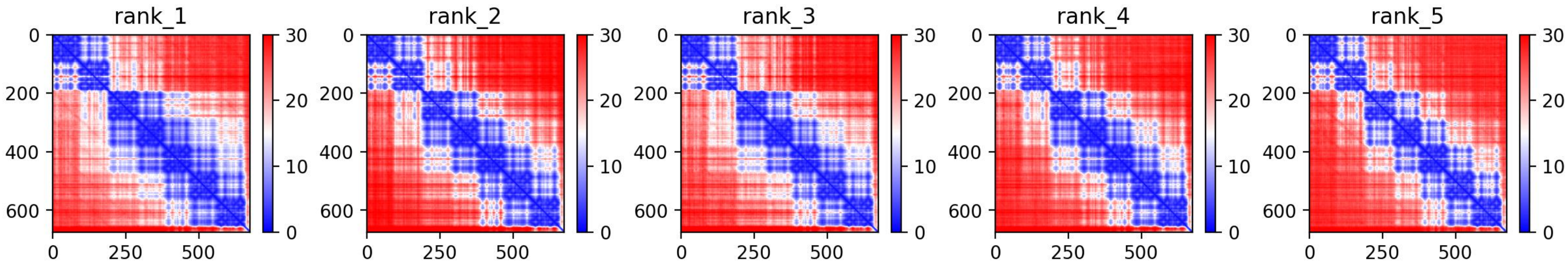

rank_1
rank_2
rank_3
rank_4
rank_5

**FMS**
**(PDGFR)**

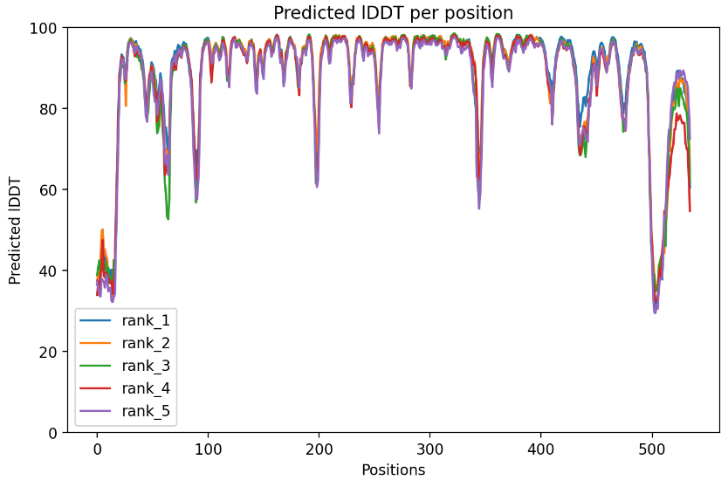

Predicted lDDT per position
Predicted lDDT
Positions
rank_1
rank_2
rank_3
rank_4
rank_5
0
20
40
60
80
100
0
100
200
300
400
500


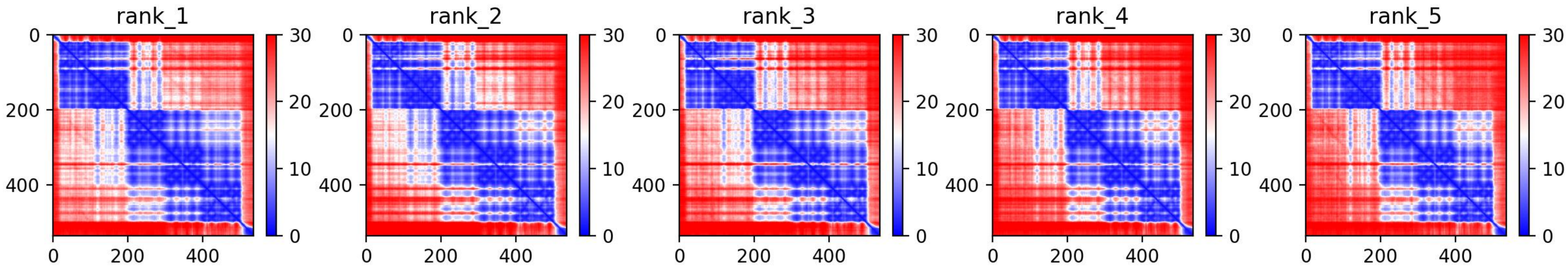

rank_1
rank_2
rank_3
rank_4
rank_5
0
200
400
0
10
20
30

**PDGFRa**
**(PDGFR)**

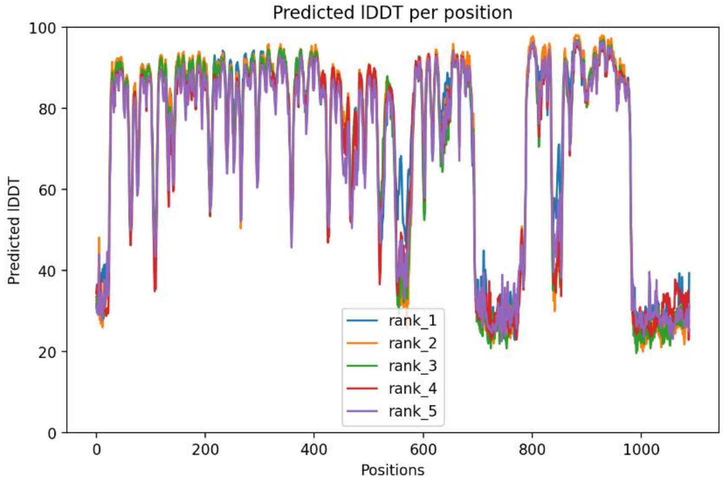


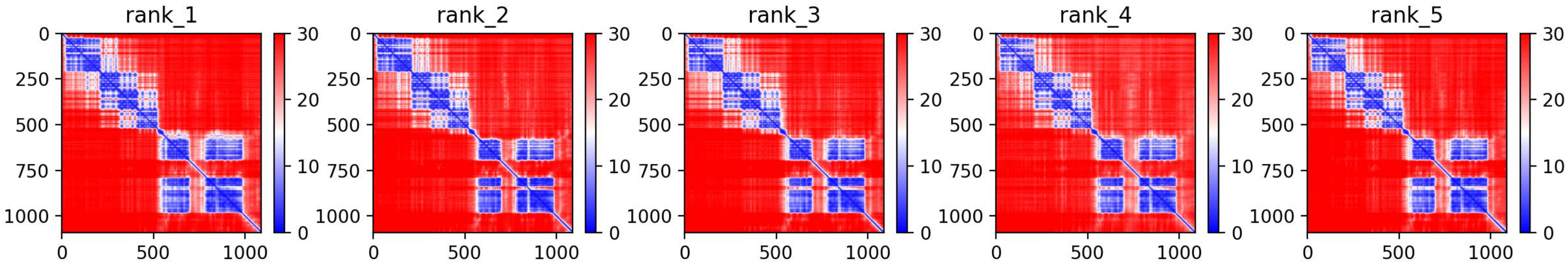

**PDGFRb (PDGFR)**

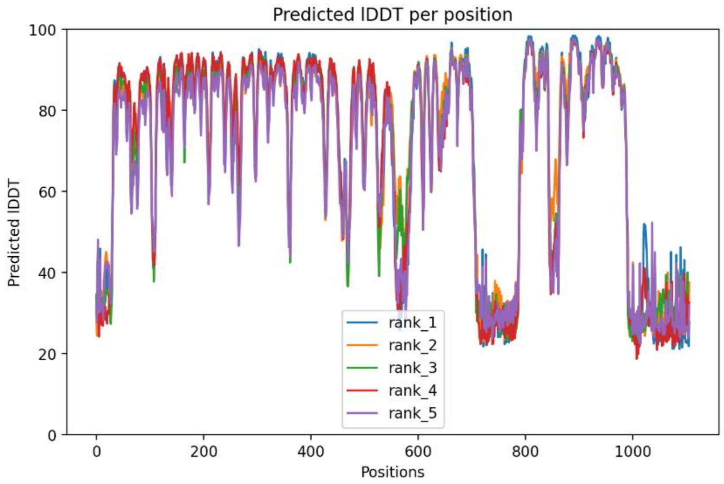


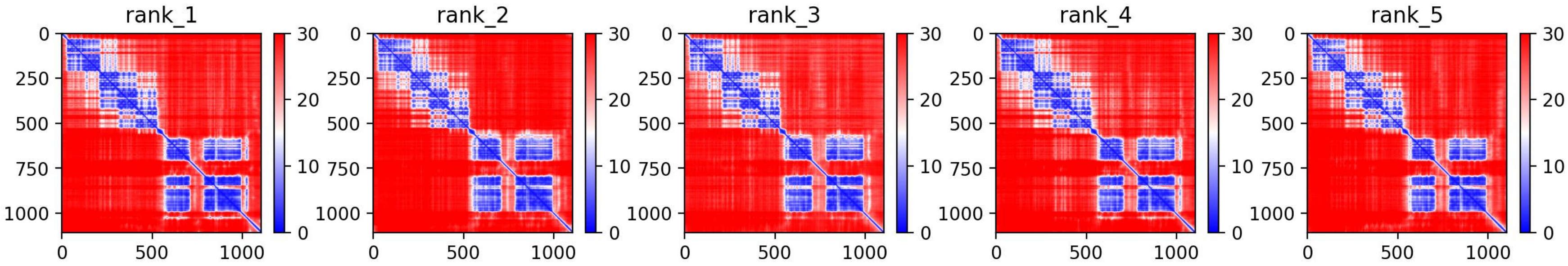

**FGFR1**
**(FGFR)**

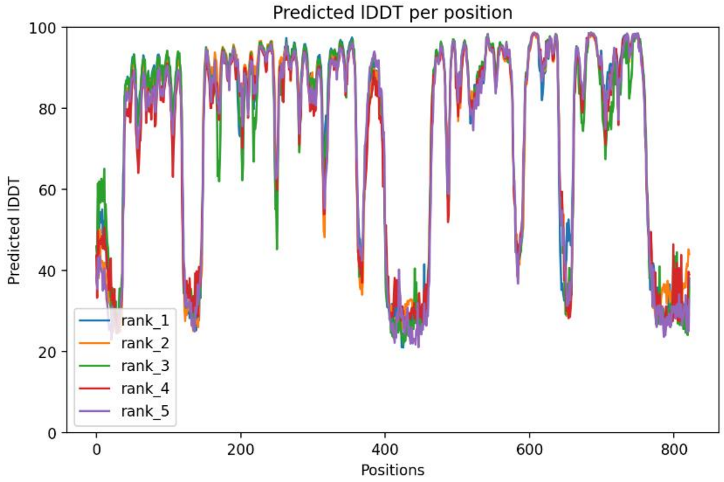

Predicted lDDT per position
Predicted lDDT
Positions
rank_1
rank_2
rank_3
rank_4
rank_5
0
20
40
60
80
100
0
200
400
600
800


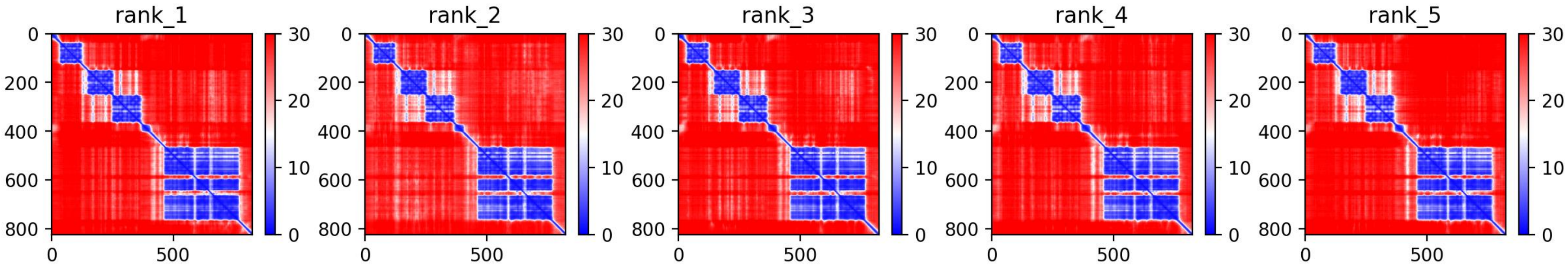

rank_1
rank_2
rank_3
rank_4
rank_5
0
200
400
600
800
0
500
30
20
10
0

**FGFR2**
**(FGFR)**

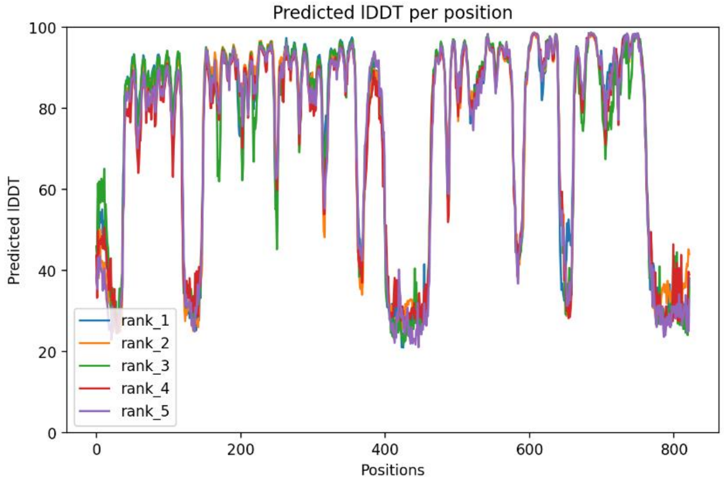

Predicted lDDT per position
Predicted lDDT
Positions
rank_1
rank_2
rank_3
rank_4
rank_5


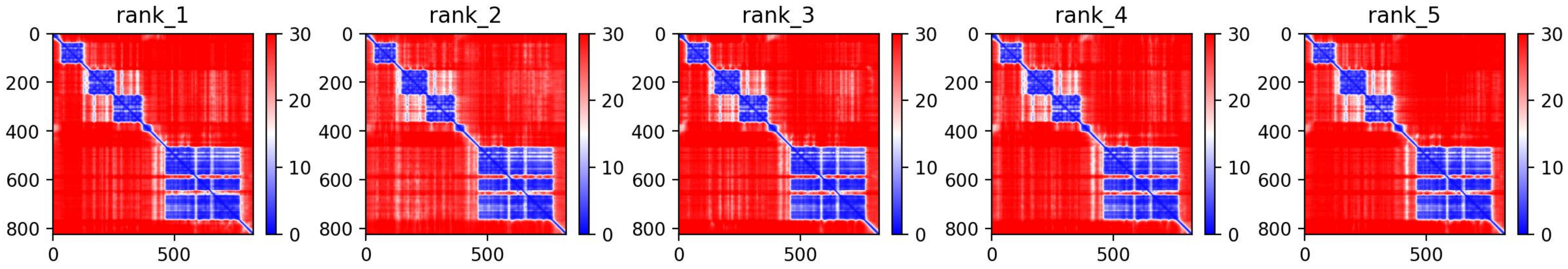

rank_1
rank_2
rank_3
rank_4
rank_5

**FGFR3**
**(FGFR)**

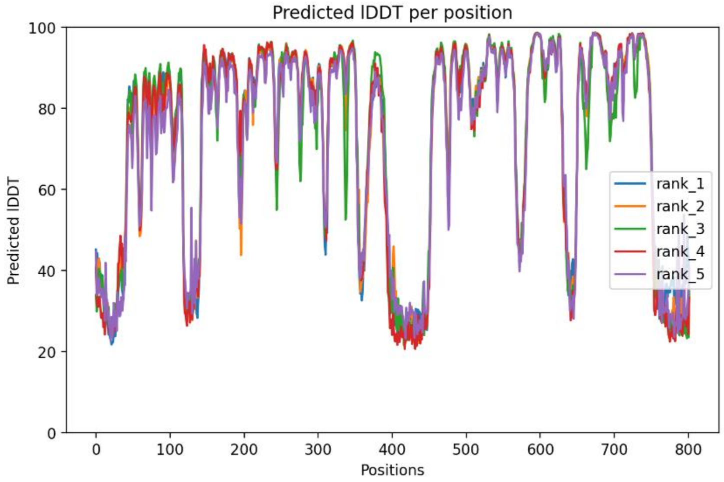
Predicted lDDT per position
Predicted lDDT
Positions
rank_1
rank_2
rank_3
rank_4
rank_5

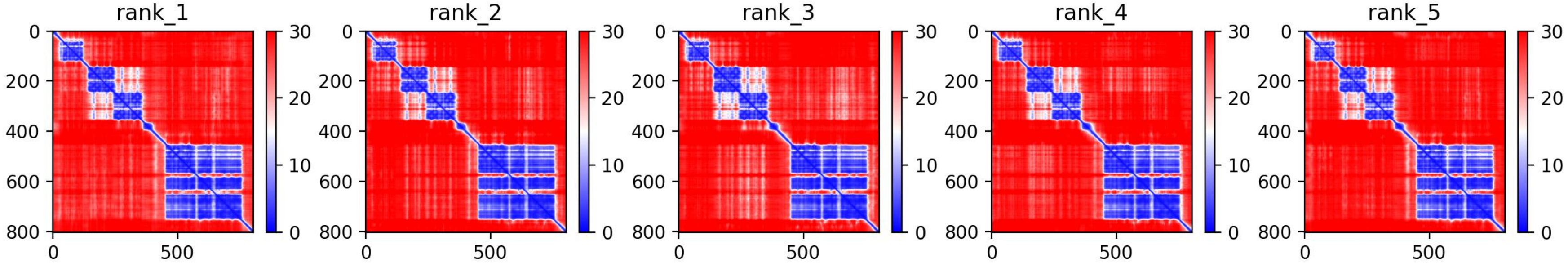
rank_1
rank_2
rank_3
rank_4
rank_5

**FGFR4**
**(FGFR)**

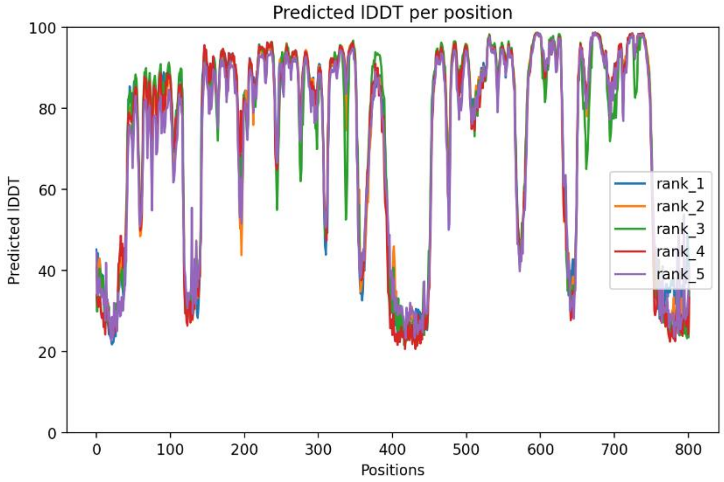

Predicted lDDT per position
Predicted lDDT
Positions
rank_1
rank_2
rank_3
rank_4
rank_5


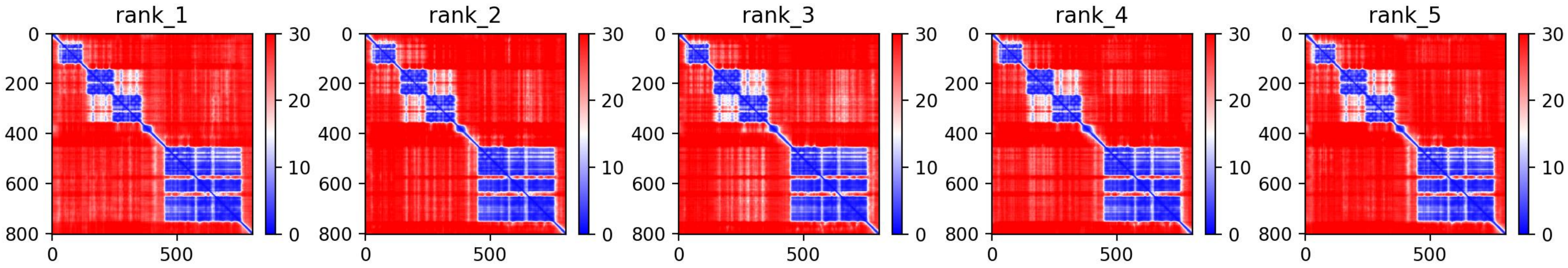

rank_1
rank_2
rank_3
rank_4
rank_5

**PTK7**
**(PTK7)**

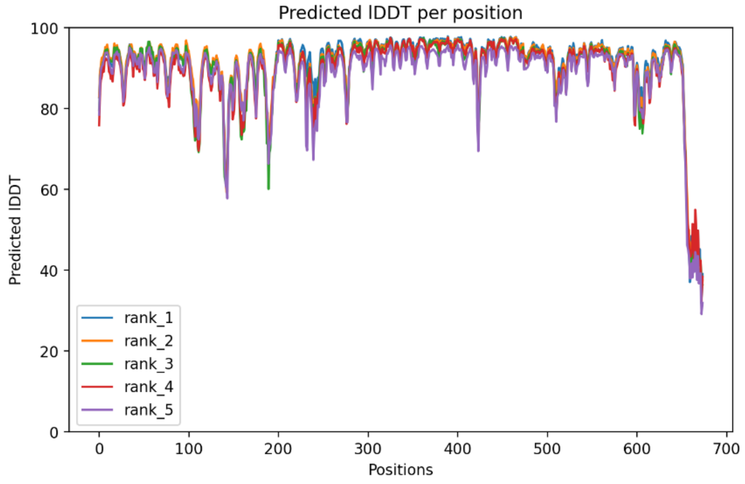

Predicted lDDT per position
Predicted lDDT
Positions
rank_1
rank_2
rank_3
rank_4
rank_5


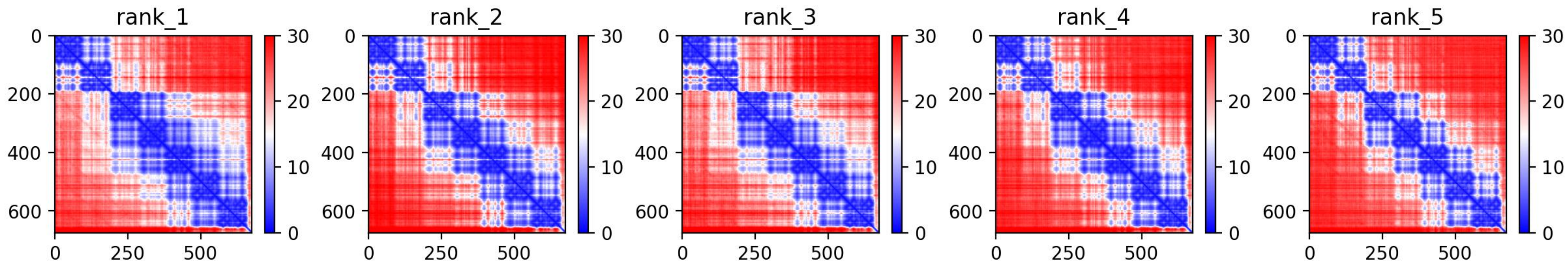

rank_1
rank_2
rank_3
rank_4
rank_5

**TrkB**
**(Trk)**

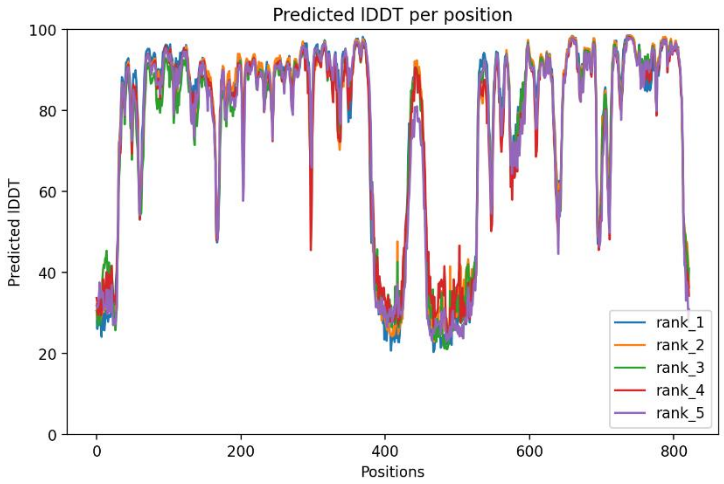

Predicted lDDT per position
Predicted lDDT
Positions
100
80
60
40
20
0
0
200
400
600
800
rank_1
rank_2
rank_3
rank_4
rank_5


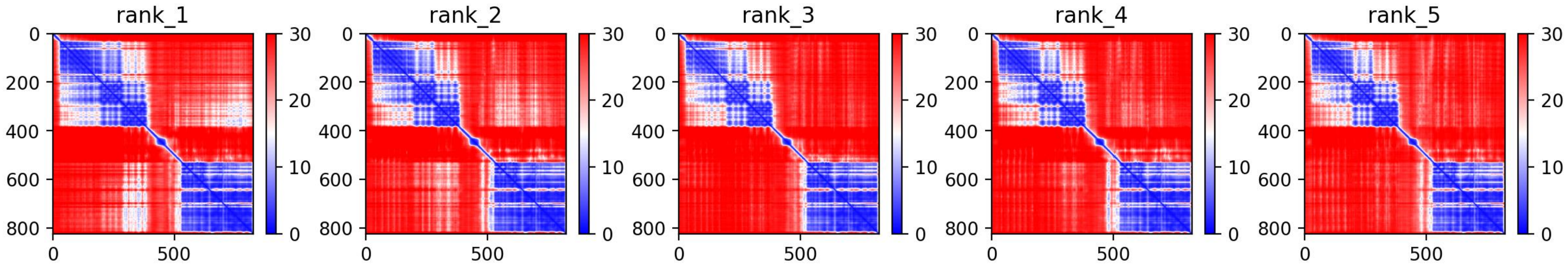

rank_1
rank_2
rank_3
rank_4
rank_5
0
200
400
600
800
0
500
30
20
10
0

**TrkC**
**(Trk)**

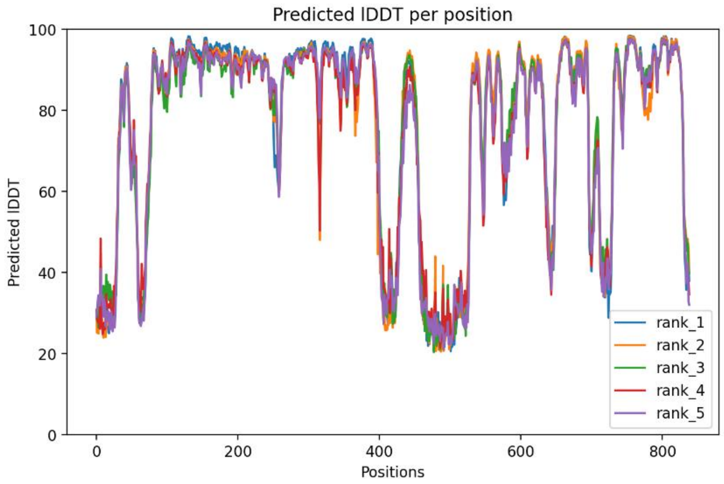
Predicted lDDT per position
Predicted lDDT
Positions
rank_1
rank_2
rank_3
rank_4
rank_5

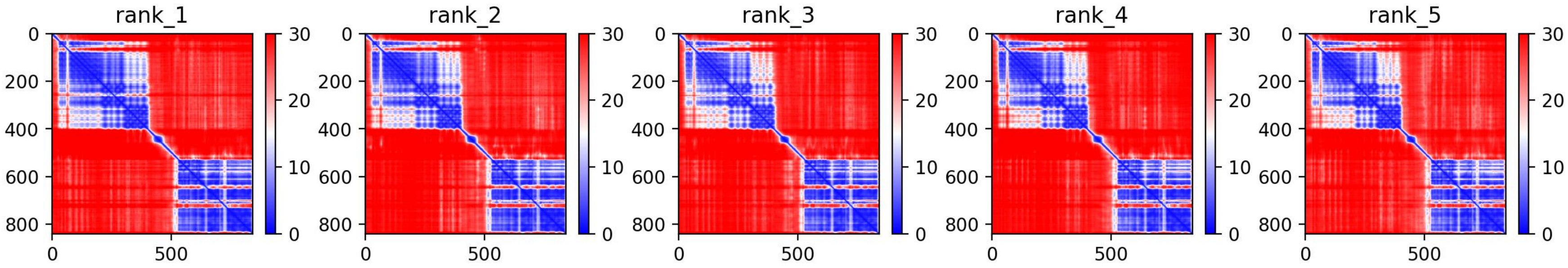
rank_1
rank_2
rank_3
rank_4
rank_5

**ROR1**
**(ROR)**

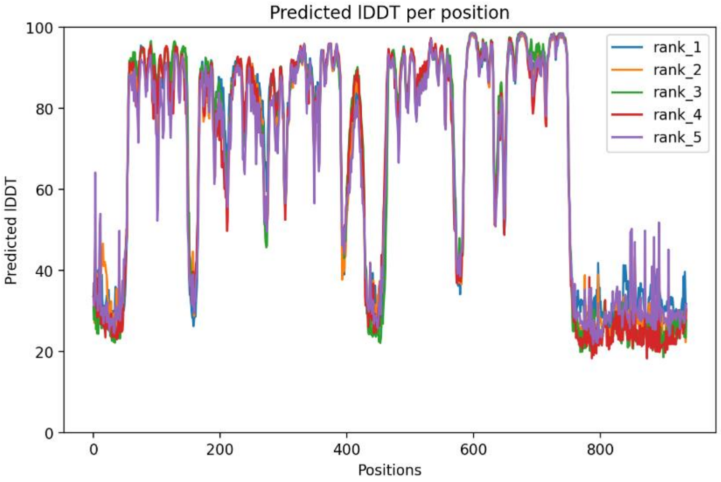

Predicted lDDT per position
Predicted lDDT
Positions
rank_1
rank_2
rank_3
rank_4
rank_5
0
20
40
60
80
100
0
200
400
600
800


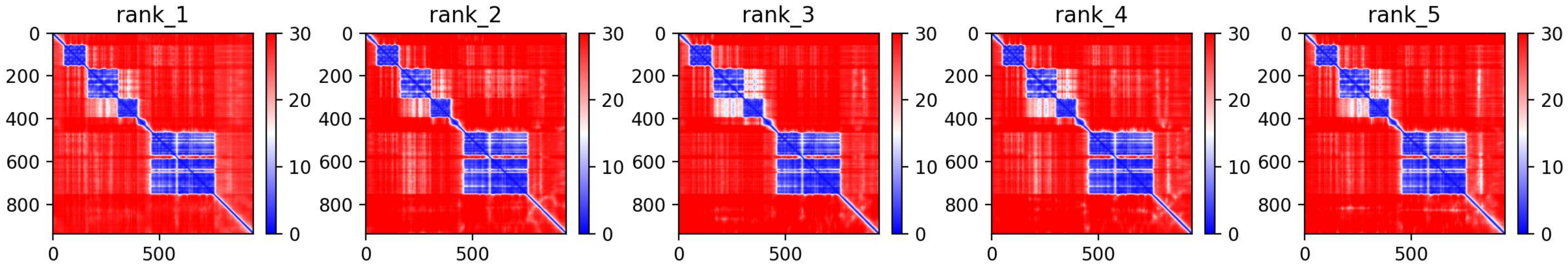

rank_1
rank_2
rank_3
rank_4
rank_5
0
200
400
600
800
0
500
30
20
10
0

**ROR2**
**(ROR)**

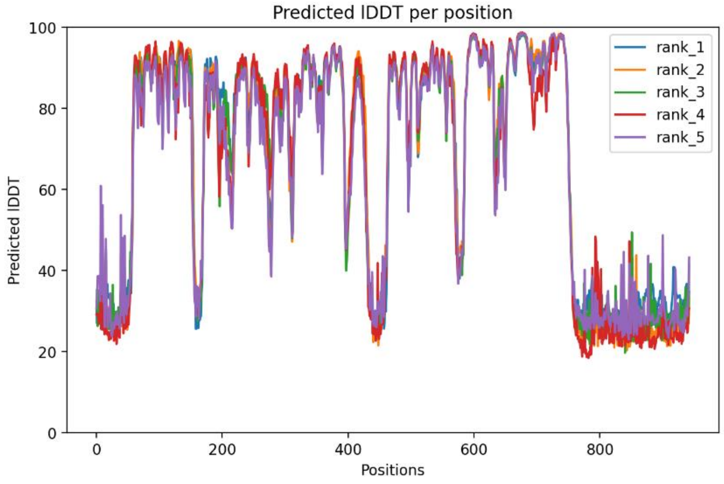

Predicted lDDT per position
Predicted lDDT
Positions
rank_1
rank_2
rank_3
rank_4
rank_5


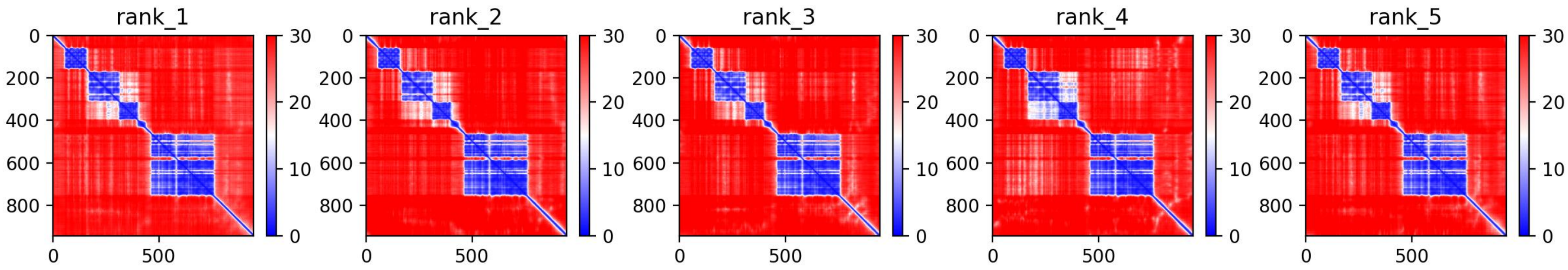

rank_1
rank_2
rank_3
rank_4
rank_5

**MuSK**
**(MuSK)**

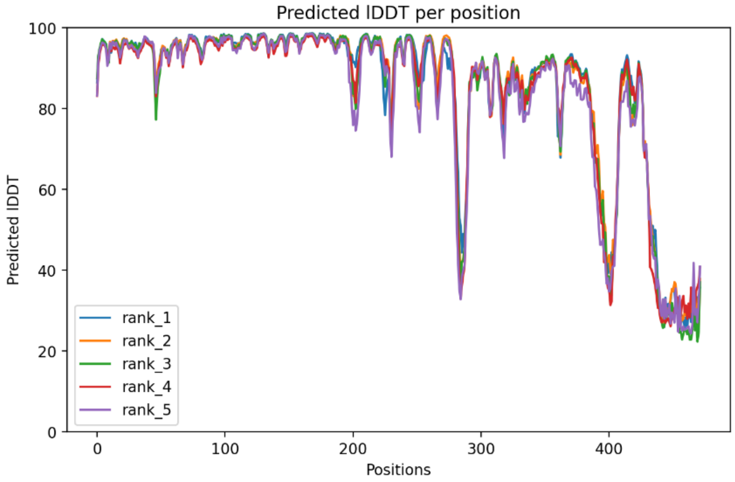

Predicted lDDT per position
Predicted lDDT
Positions
rank_1
rank_2
rank_3
rank_4
rank_5


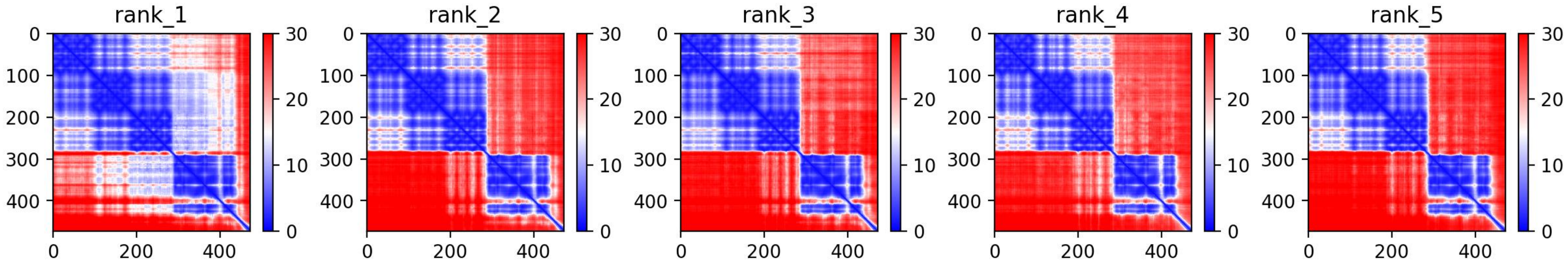

rank_1
rank_2
rank_3
rank_4
rank_5

**MET**
**(MET)**

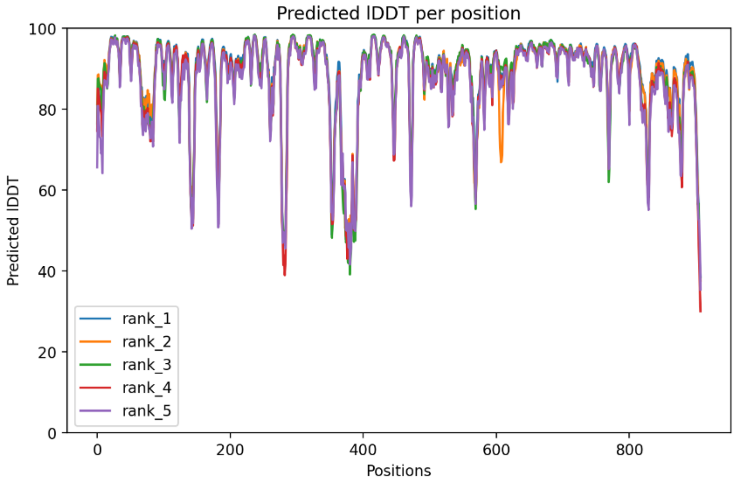
Predicted lDDT per position
Predicted lDDT
Positions
rank_1
rank_2
rank_3
rank_4
rank_5
100
80
60
40
20
0
0
200
400
600
800

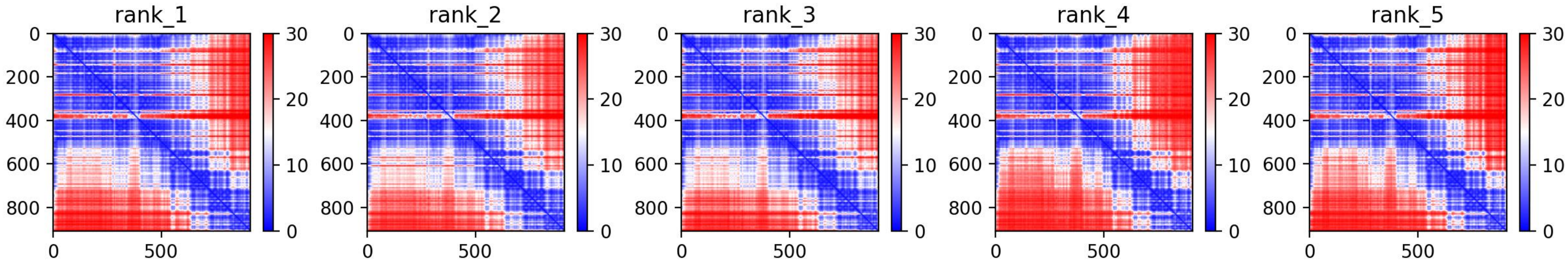
rank_1
rank_2
rank_3
rank_4
rank_5
0
200
400
600
800
0
500
30
20
10
0

**RON**
**(MET)**

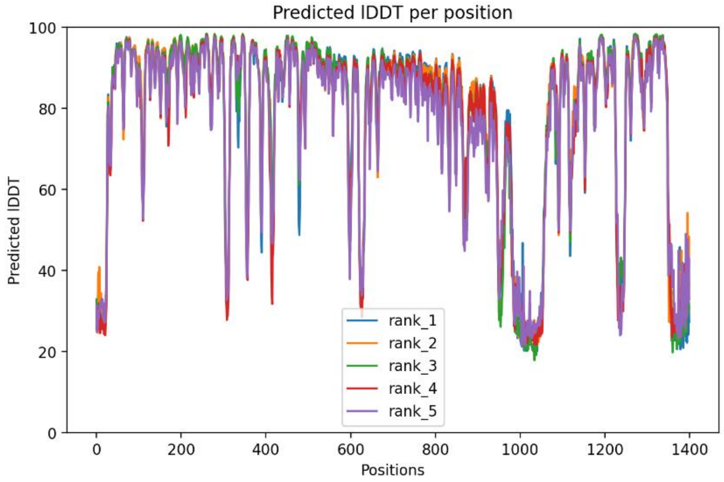
Predicted lDDT per position
Predicted lDDT
Positions
rank_1
rank_2
rank_3
rank_4
rank_5

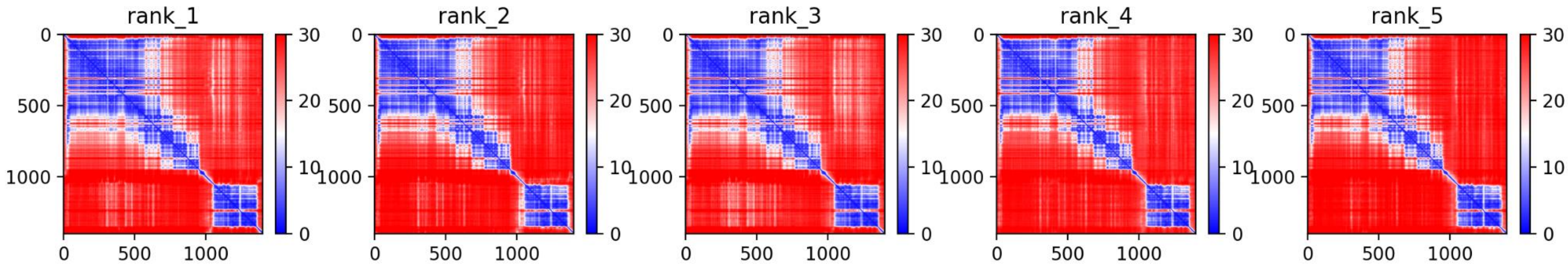
rank_1
rank_2
rank_3
rank_4
rank_5

**Axl**
**(Axl)**

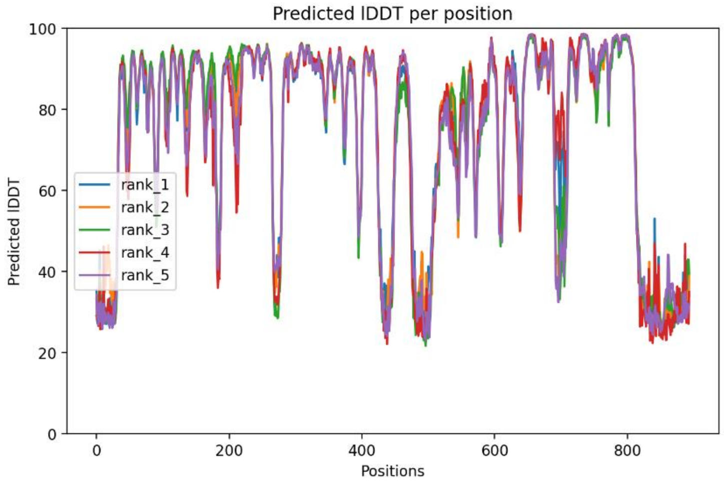
Predicted lDDT per position
Predicted lDDT
Positions
rank_1
rank_2
rank_3
rank_4
rank_5
100
80
60
40
20
0
0
200
400
600
800

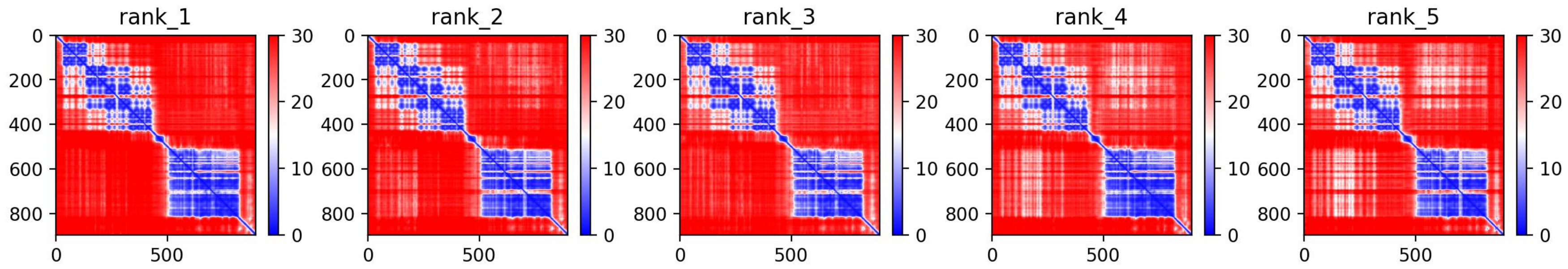
rank_1
rank_2
rank_3
rank_4
rank_5
0
200
400
600
800
0
500
30
20
10
0

**MER**
**(Axl)**

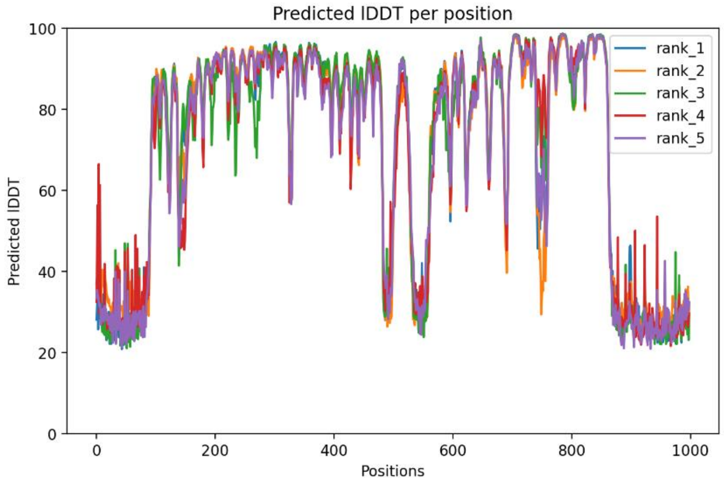
Predicted lDDT per position
Predicted lDDT
Positions
100
80
60
40
20
0
0
200
400
600
800
1000
rank_1
rank_2
rank_3
rank_4
rank_5

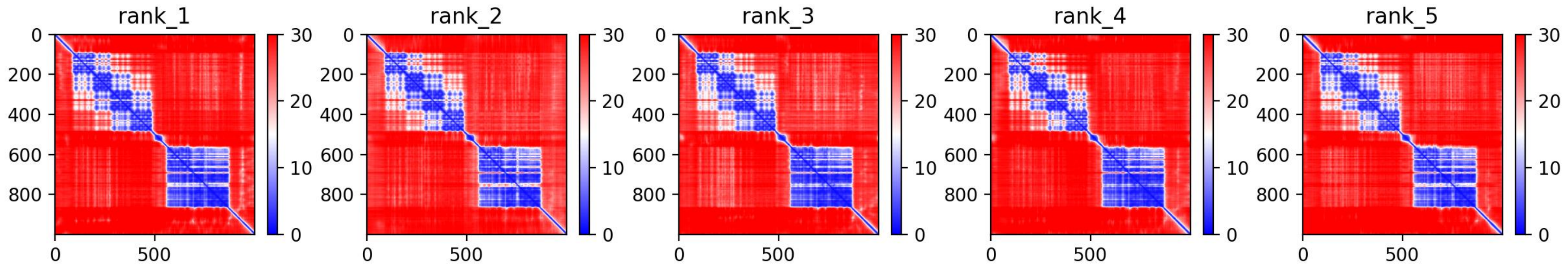
rank_1
rank_2
rank_3
rank_4
rank_5
0
200
400
600
800
0
500
30
20
10
0

**Tyro3 (Axl)**

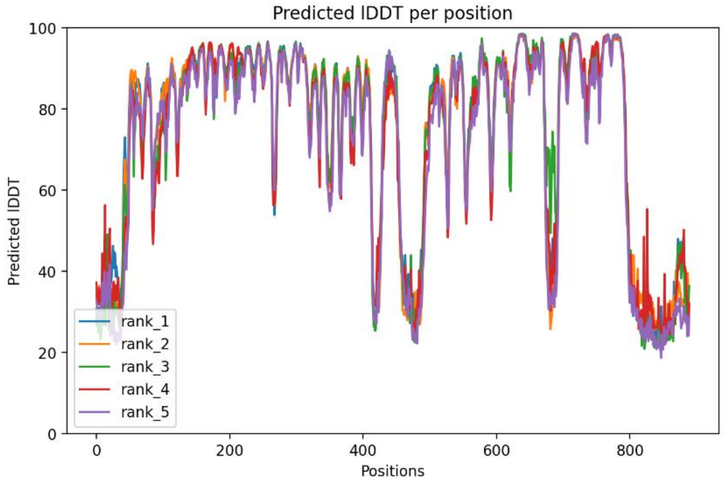

Predicted lDDT per position
Predicted lDDT
Positions
rank_1
rank_2
rank_3
rank_4
rank_5


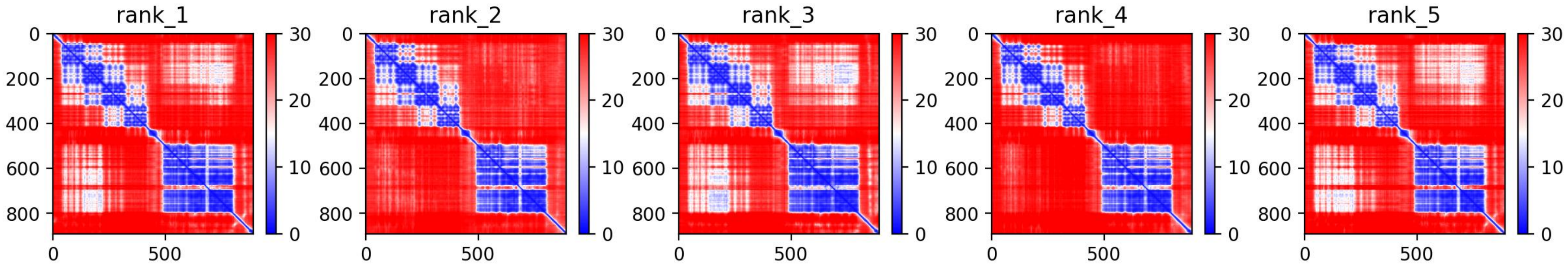

rank_1
rank_2
rank_3
rank_4
rank_5

**Tie1**
**(Tie)**

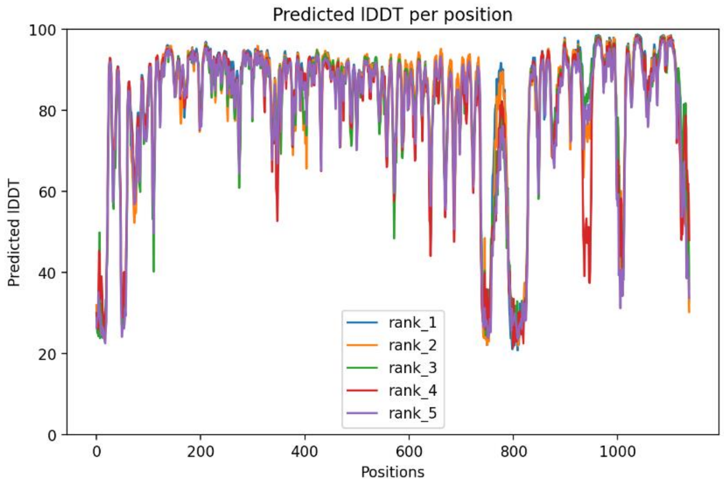
Predicted lDDT per position
Predicted lDDT
Positions
rank_1
rank_2
rank_3
rank_4
rank_5

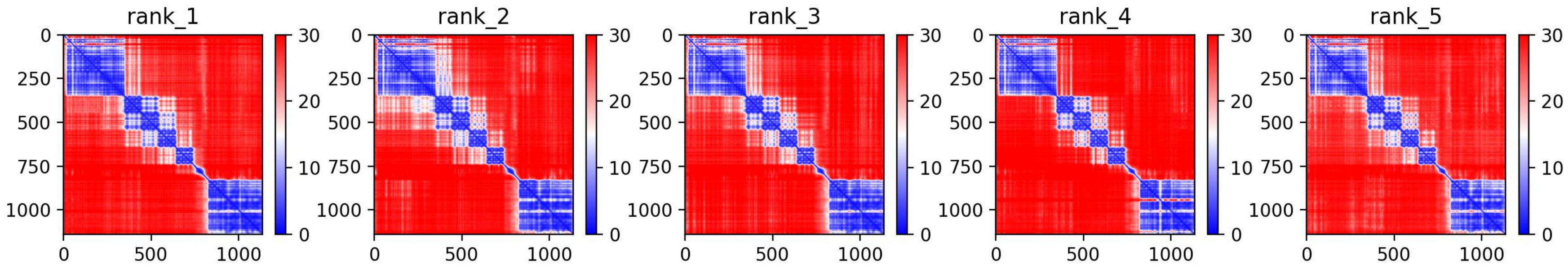
rank_1
rank_2
rank_3
rank_4
rank_5

**EphR1**
**(EphR)**

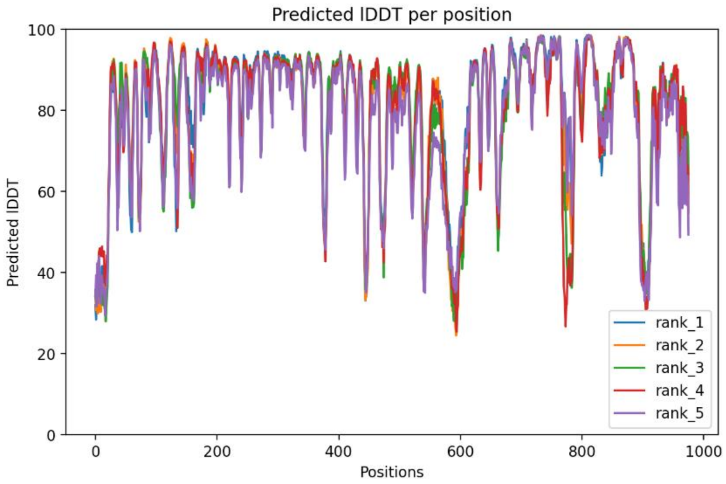

Predicted lDDT per position
Predicted lDDT
Positions
100
80
60
40
20
0
0
200
400
600
800
1000
rank_1
rank_2
rank_3
rank_4
rank_5


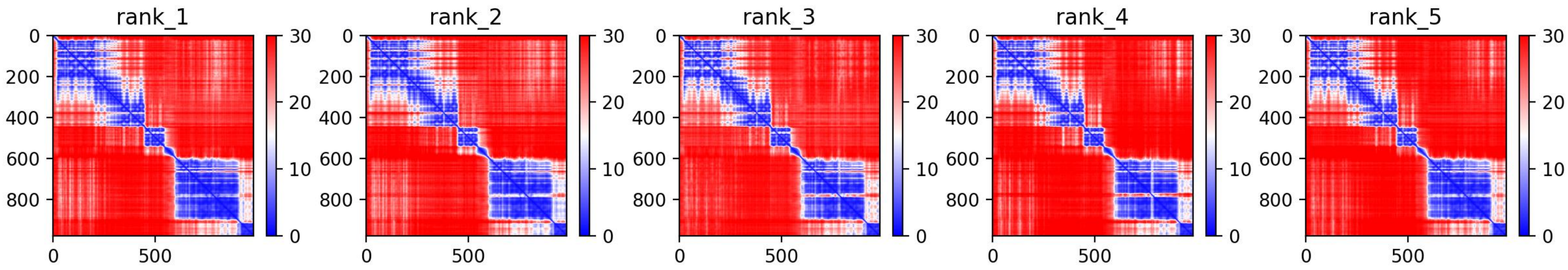

rank_1
rank_2
rank_3
rank_4
rank_5
0
200
400
600
800
0
500
30
20
10
0

**EphR3**
**(EphR)**

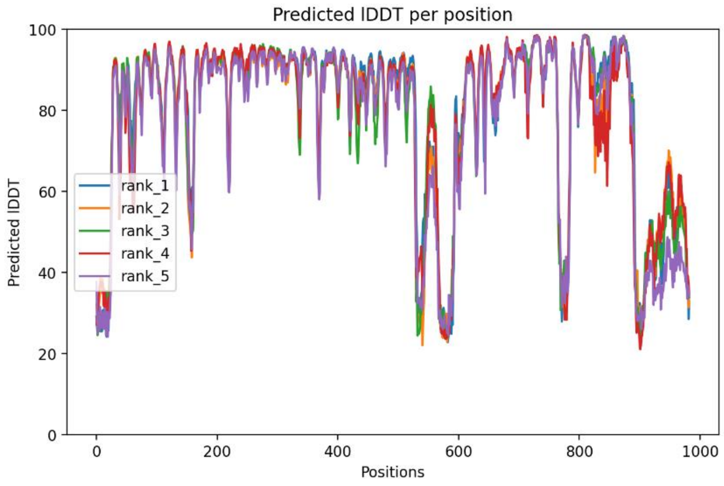
Predicted lDDT per position
Predicted lDDT
Positions
rank_1
rank_2
rank_3
rank_4
rank_5

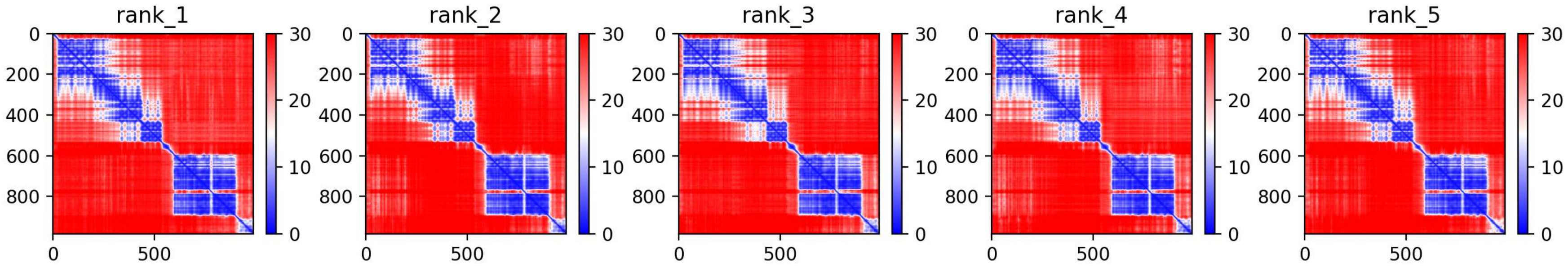
rank_1
rank_2
rank_3
rank_4
rank_5

**EphR5**
**(EphR)**

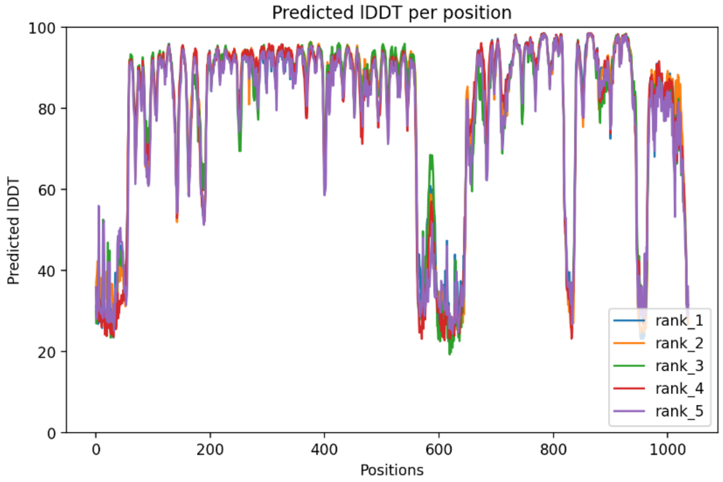
Predicted lDDT per position
Predicted lDDT
Positions
0
20
40
60
80
100
0
200
400
600
800
1000
rank_1
rank_2
rank_3
rank_4
rank_5

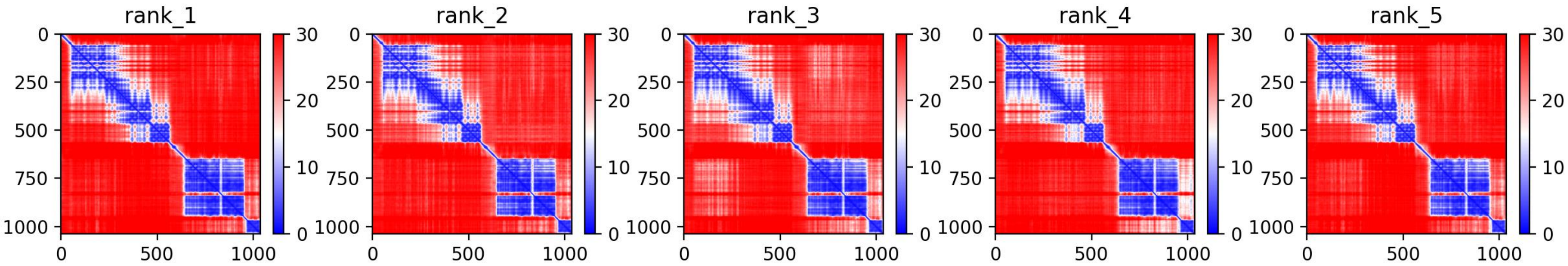
rank_1
rank_2
rank_3
rank_4
rank_5
0
250
500
750
1000
0
500
1000
0
10
20
30

**EphR6 (EphR)**

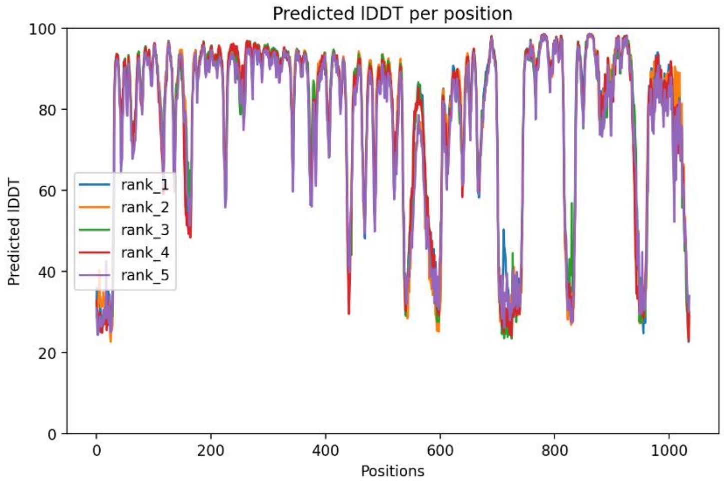
Predicted lDDT per position
Predicted lDDT
Positions
rank_1
rank_2
rank_3
rank_4
rank_5

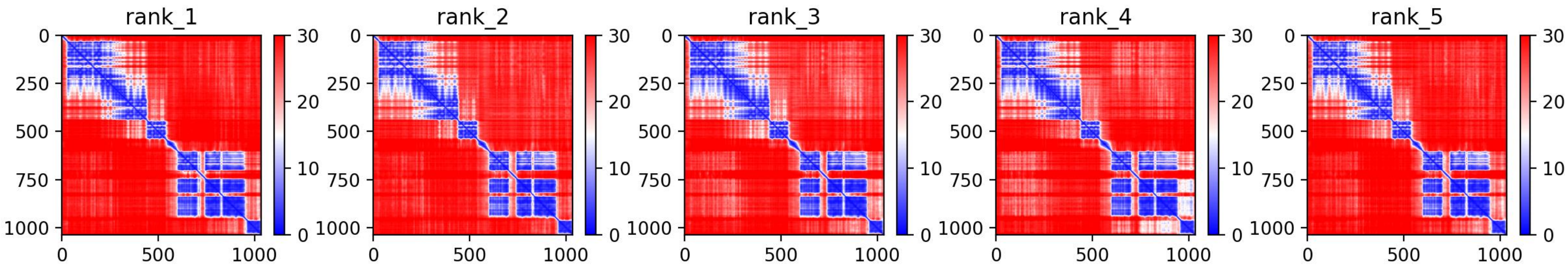
rank_1
rank_2
rank_3
rank_4
rank_5

**EphR7**
**(EphR)**

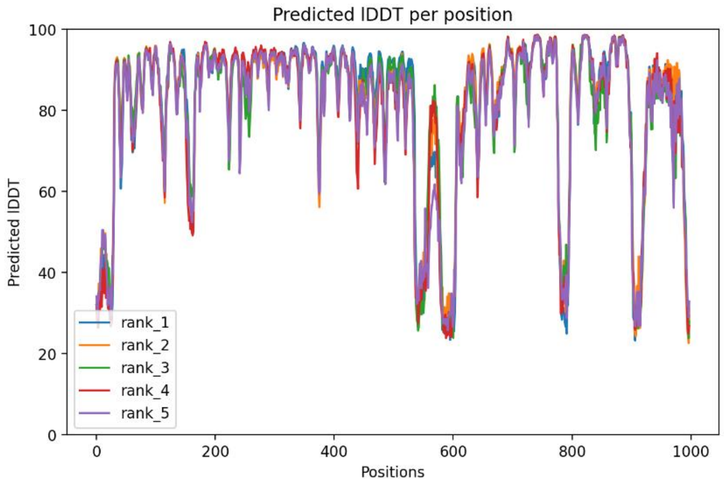
Predicted lDDT per position
Predicted lDDT
Positions
rank_1
rank_2
rank_3
rank_4
rank_5
0
20
40
60
80
100
0
200
400
600
800
1000

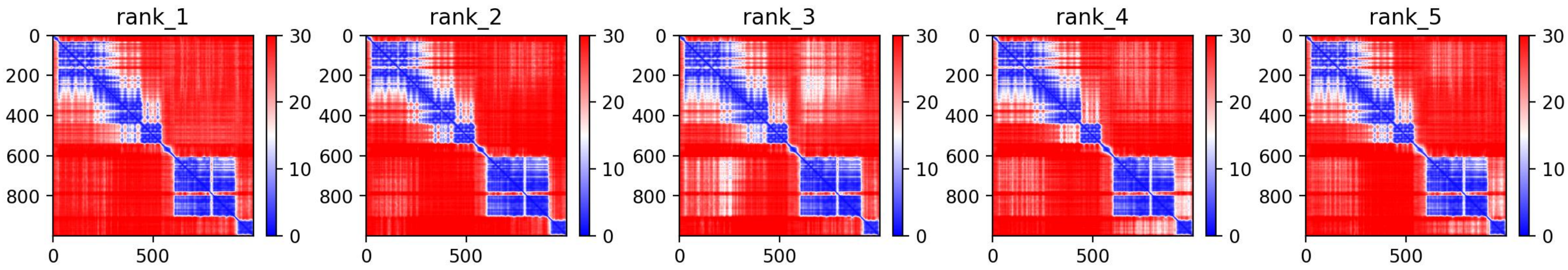
rank_1
rank_2
rank_3
rank_4
rank_5
0
200
400
600
800
0
500
30
20
10
0

**EphRA8**
**(EphR)**

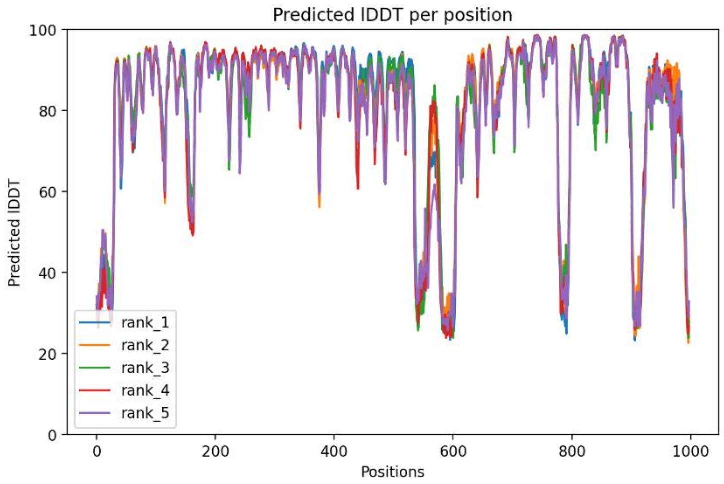
Predicted lDDT per position
Predicted lDDT
Positions
rank_1
rank_2
rank_3
rank_4
rank_5

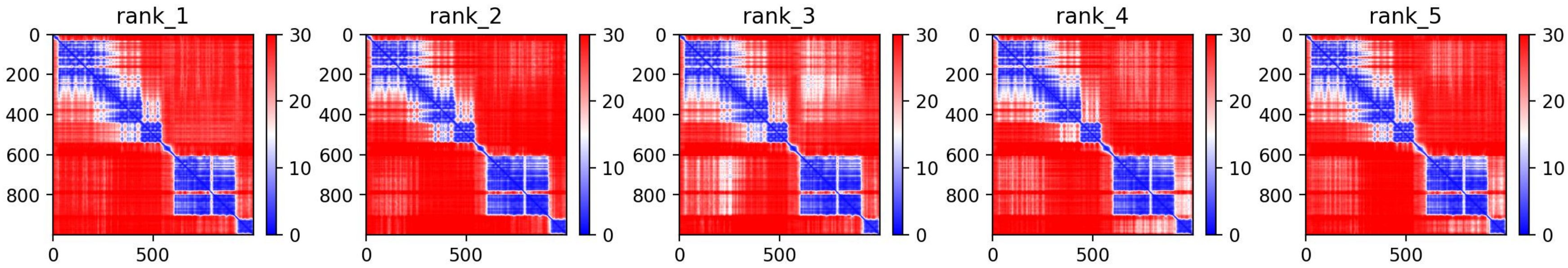
rank_1
rank_2
rank_3
rank_4
rank_5

**EphRA10 (EphR)**

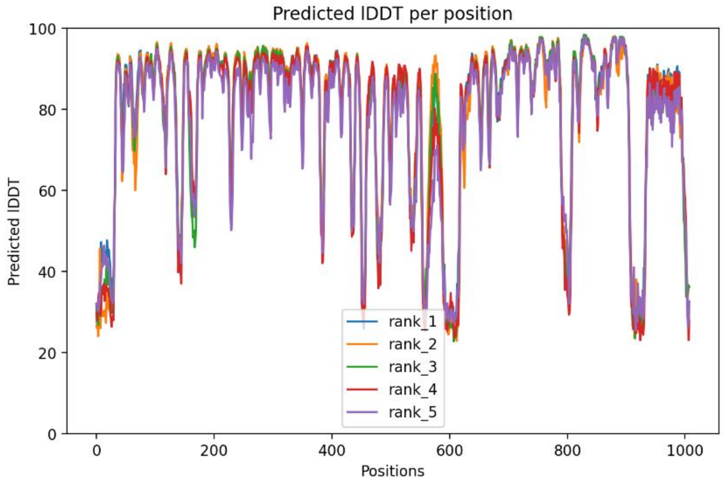


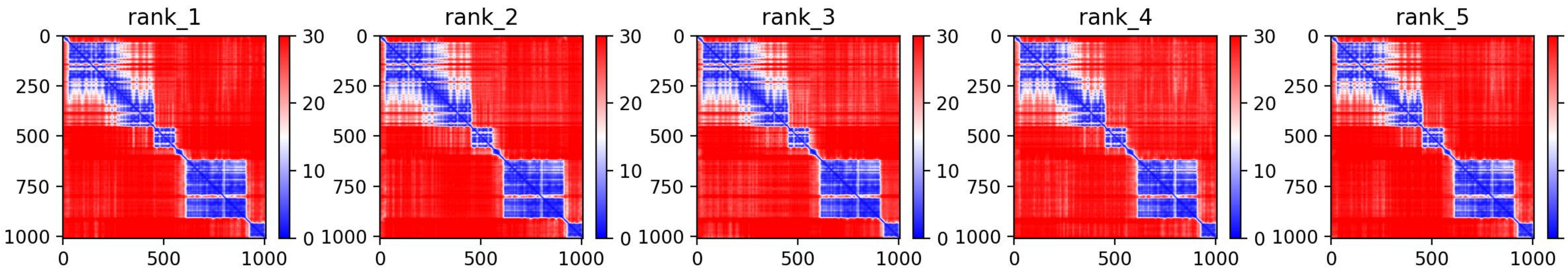

**EphRB1**
**(EphR)**

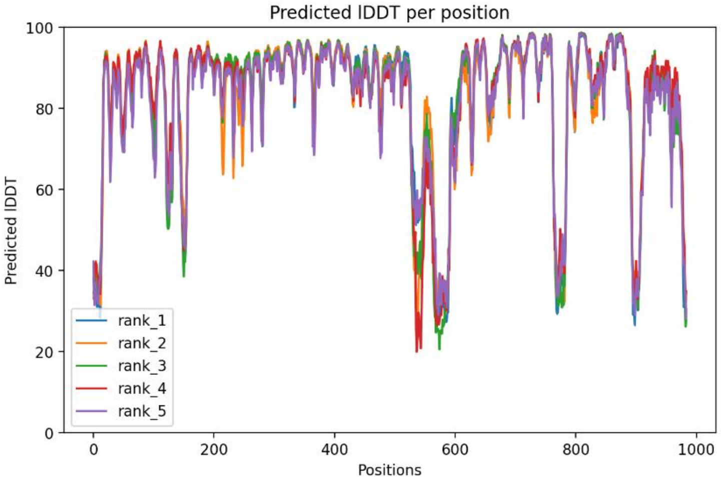
Predicted lDDT per position
Predicted lDDT
Positions
rank_1
rank_2
rank_3
rank_4
rank_5


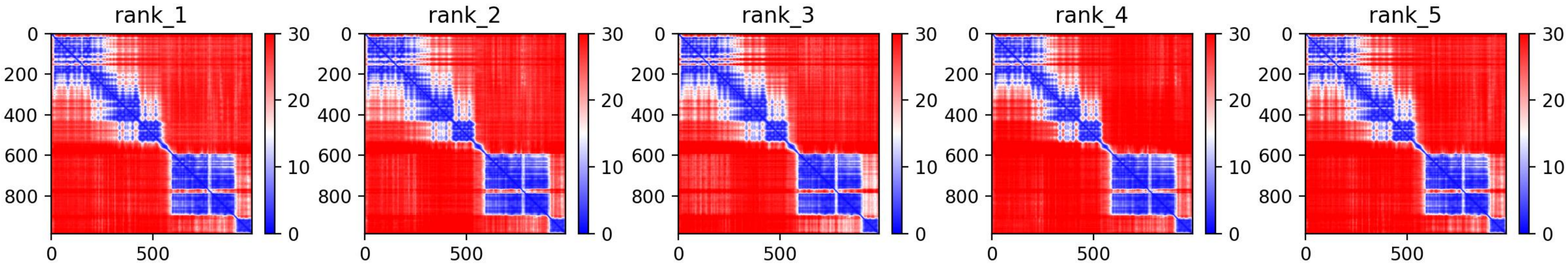
rank_1
rank_2
rank_3
rank_4
rank_5

**EphRB3**
**(EphR)**

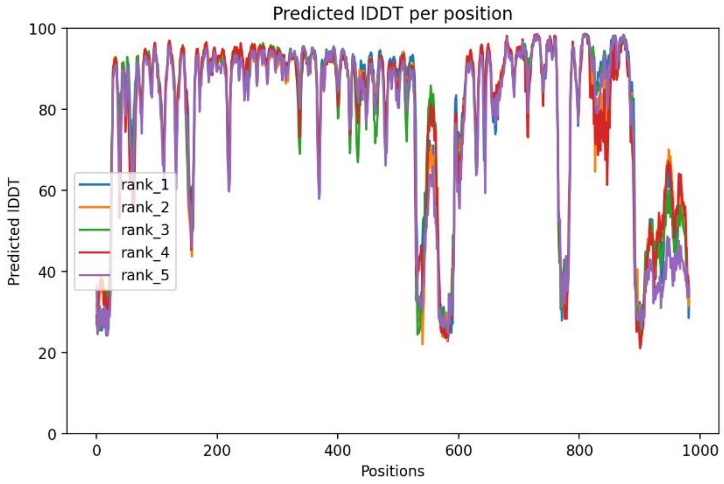
Predicted lDDT per position
Predicted lDDT
Positions
rank_1
rank_2
rank_3
rank_4
rank_5

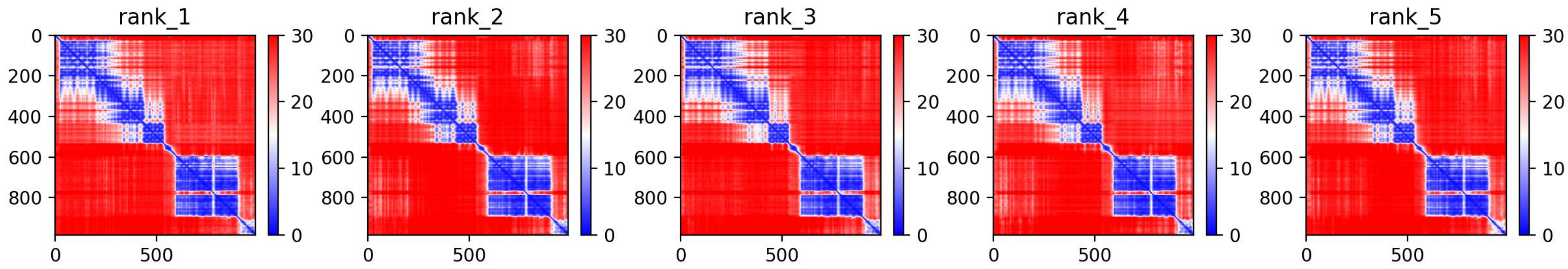
rank_1
rank_2
rank_3
rank_4
rank_5

**EphRB4**
**(EphR)**

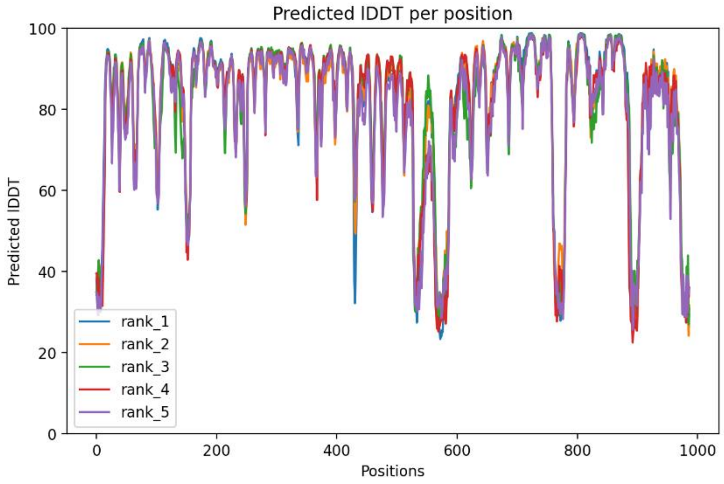
Predicted lDDT per position
Predicted lDDT
Positions
rank_1
rank_2
rank_3
rank_4
rank_5
0
20
40
60
80
100
0
200
400
600
800
1000

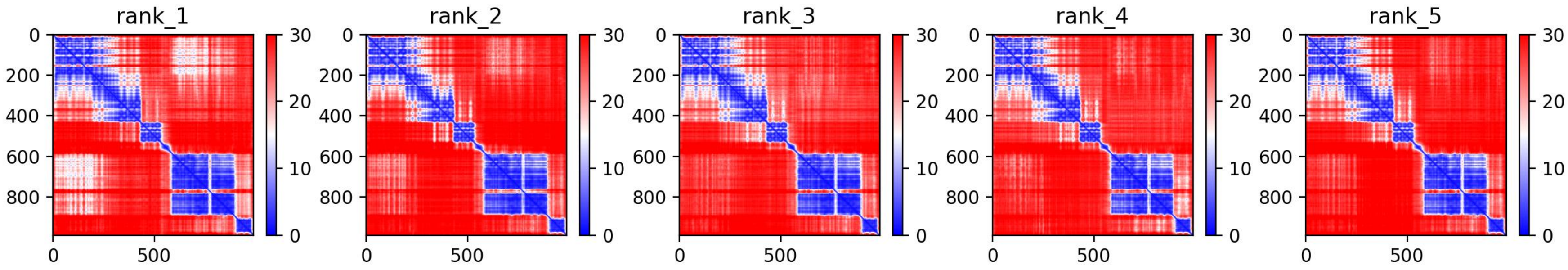
rank_1
rank_2
rank_3
rank_4
rank_5
0
200
400
600
800
0
500
0
10
20
30

**EphRB6**
**(EphR)**

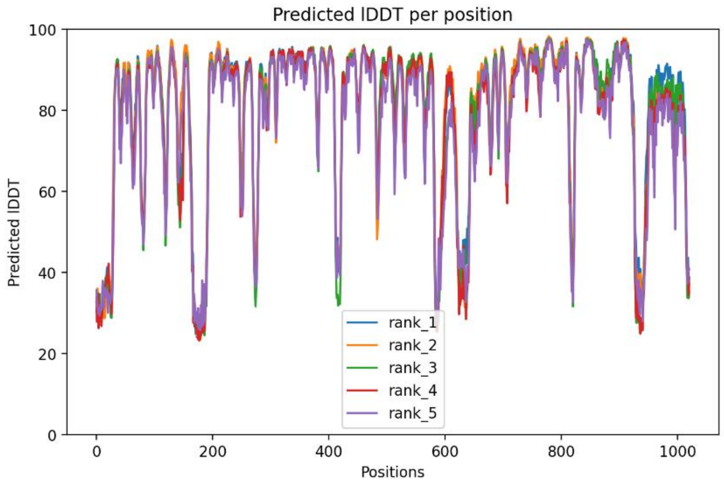
Predicted lDDT per position
Predicted lDDT
Positions
rank_1
rank_2
rank_3
rank_4
rank_5

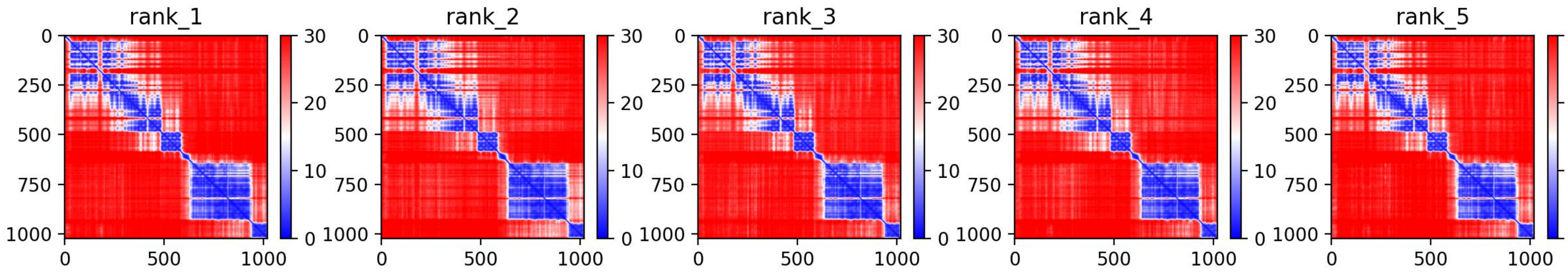
rank_1
rank_2
rank_3
rank_4
rank_5

**Ryk**
**(Ryk)**

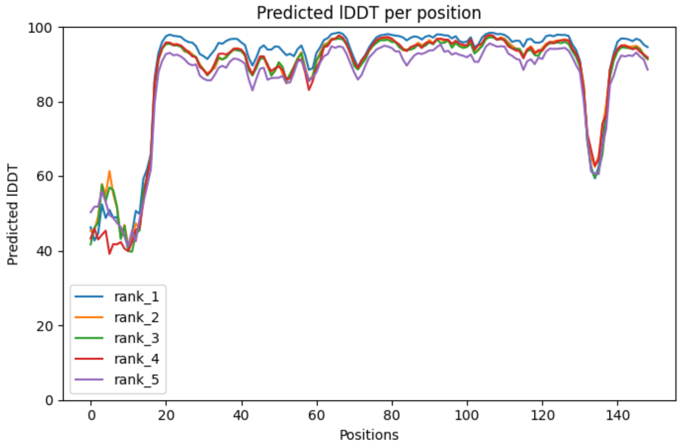
Predicted lDDT per position
Predicted lDDT
Positions
rank_1
rank_2
rank_3
rank_4
rank_5

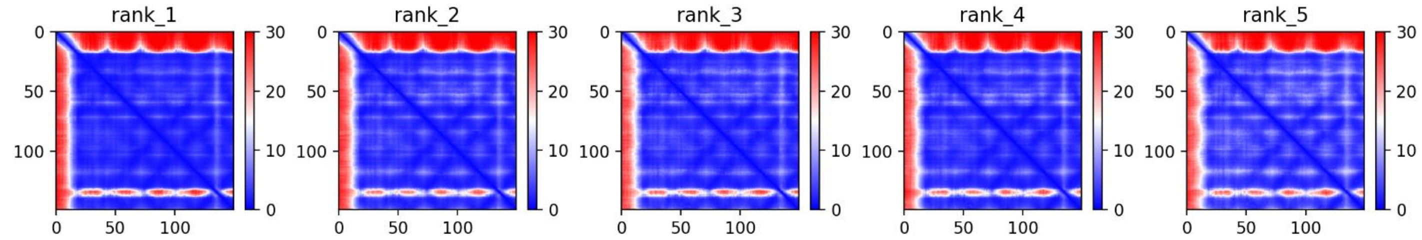
rank_1
rank_2
rank_3
rank_4
rank_5

**DDR2**
**(DDR)**

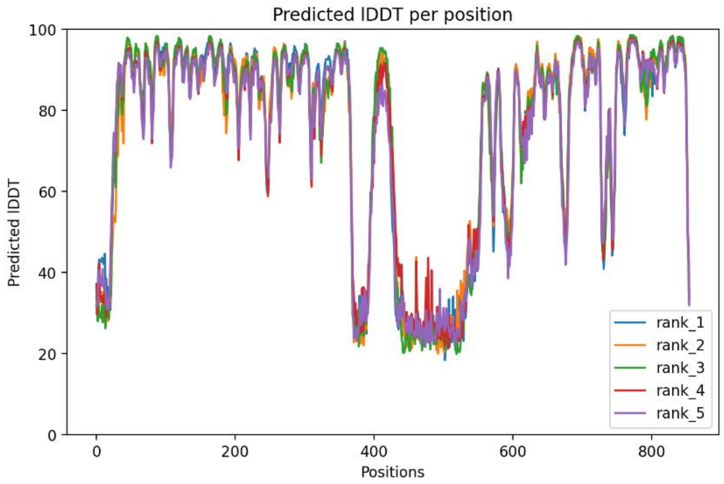
Predicted lDDT per position
Predicted lDDT
Positions
100
80
60
40
20
0
0
200
400
600
800
rank_1
rank_2
rank_3
rank_4
rank_5

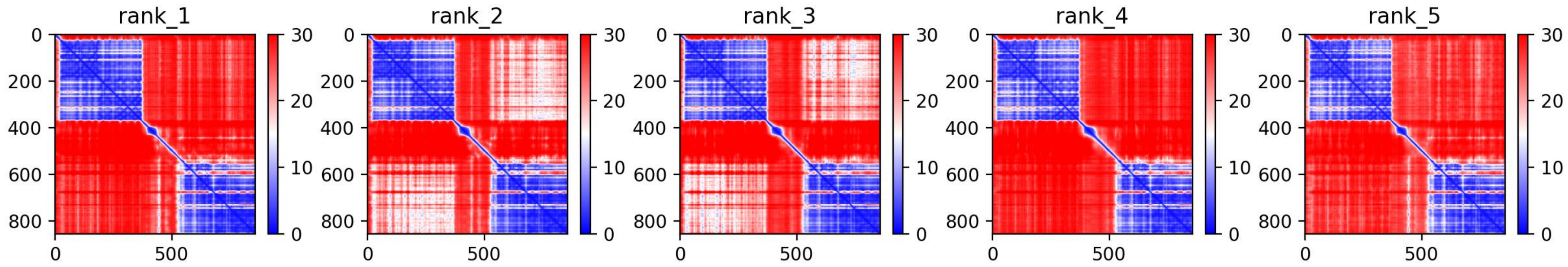
rank_1
rank_2
rank_3
rank_4
rank_5
0
200
400
600
800
0
500
30
20
10
0

**ROS**
**(ROS)**

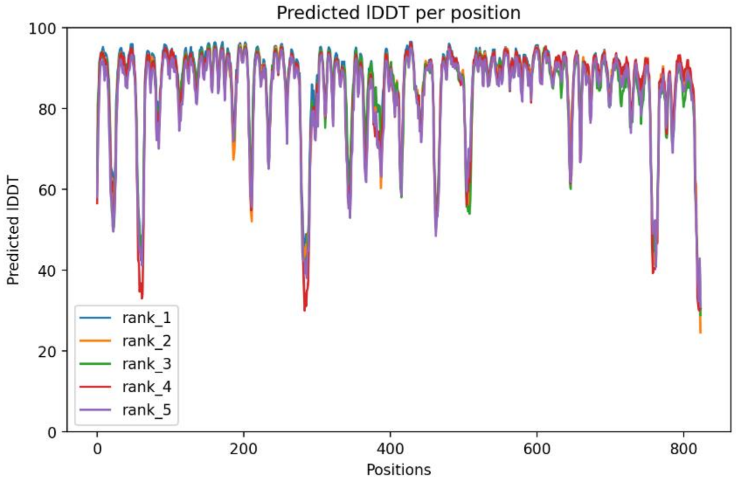

Predicted lDDT per position
Predicted lDDT
Positions
rank_1
rank_2
rank_3
rank_4
rank_5


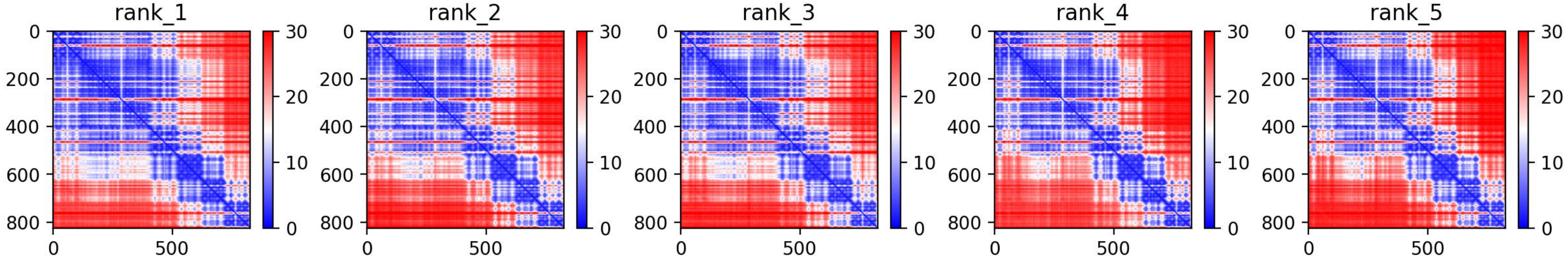

rank_1
rank_2
rank_3
rank_4
rank_5

**ALK**
**(ALK)**

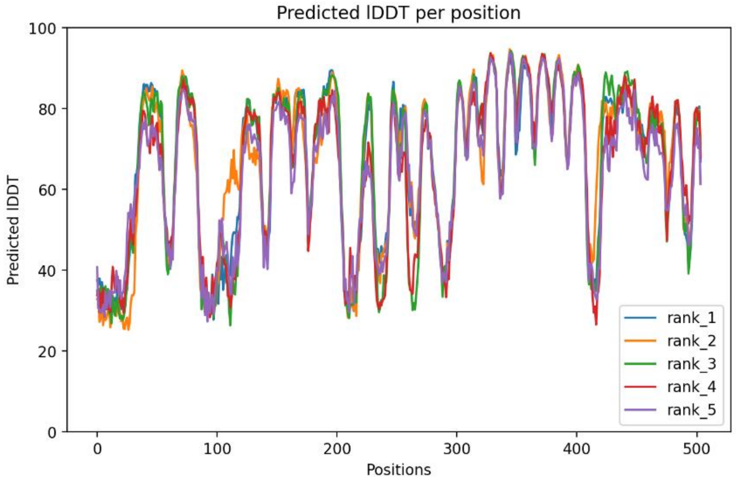

Predicted lDDT per position
Predicted lDDT
Positions
rank_1
rank_2
rank_3
rank_4
rank_5


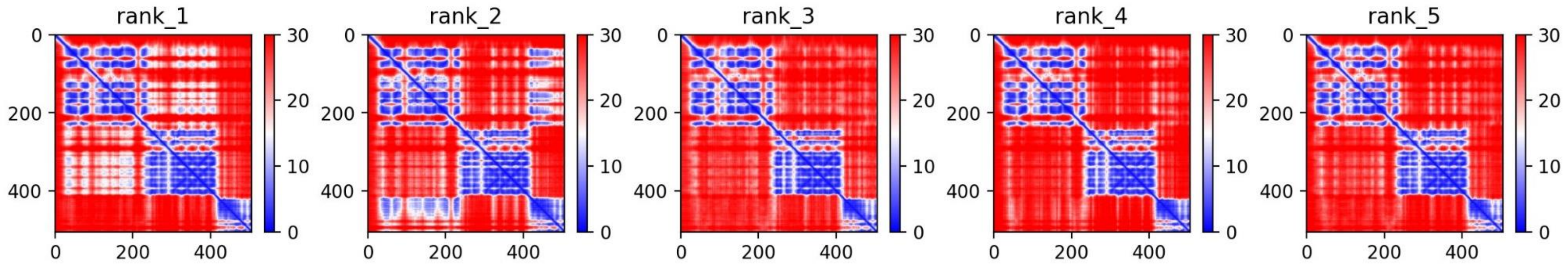

rank_1
rank_2
rank_3
rank_4
rank_5

**LTK**
**(ALK)**

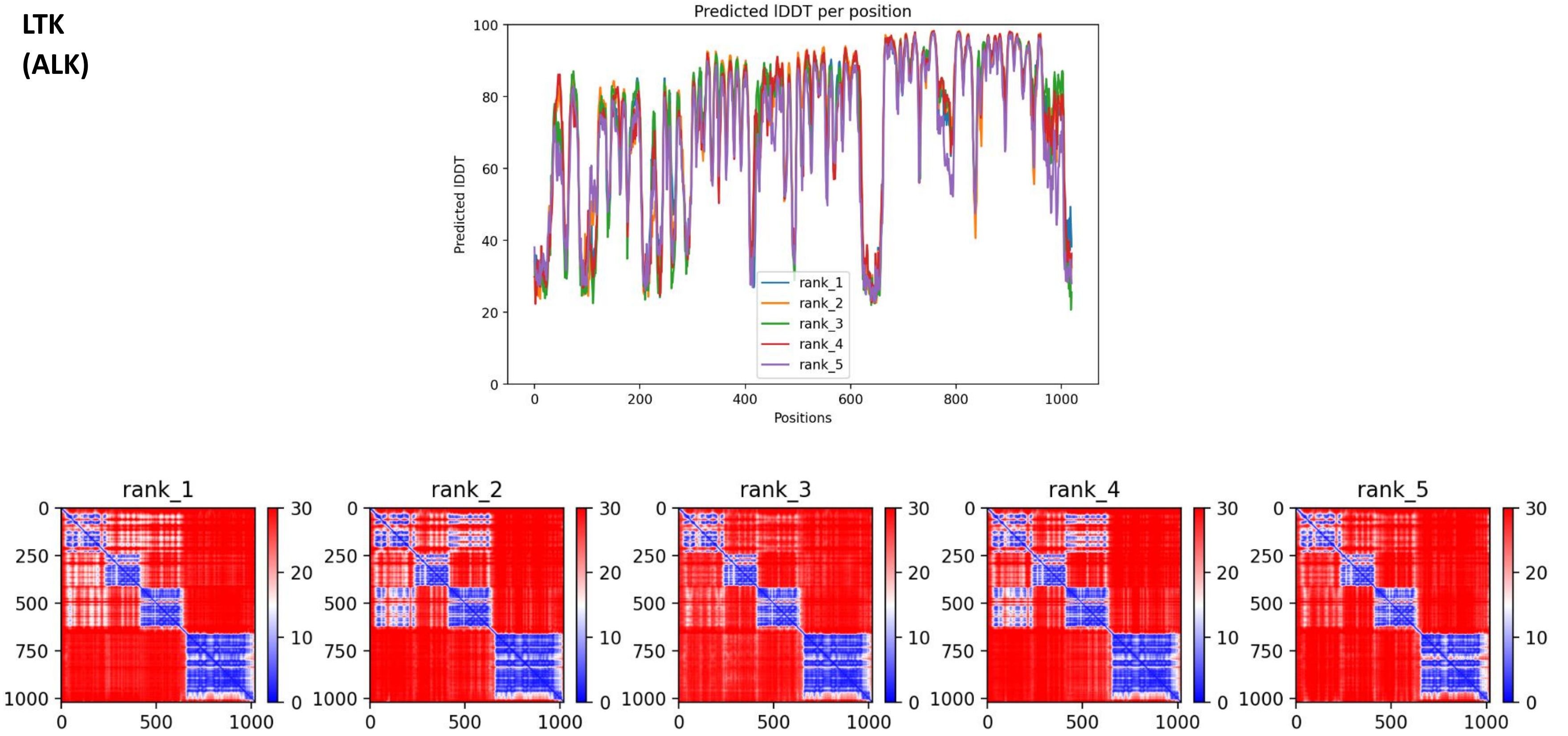


**Supplementary Figure 1. AlphaFold2 model quality measures. For each modeled RTK (with RTK family in parenthesis) are shown the pLDDT plot, assessing local prediction confidence, and the PAE plot for each model (ranked 1-5), for evaluating the prediction accuracy of inter-domain positioning.**
