## Supplementary Table 1 for "Structural conservation and divergence across the Receptor Tyrosine Kinase superfamily"

| RTK family | RTK name | UniPprot entry | Domain identifier | Functional interactions | 3D fold |
|---|---|---|---|---|---|
| PDGFR | KIT | P10721 | 2E9W-A1 | LBD | Ig-like (1) |
| | | | 2E9W-A2 | LBD | Ig-like (1) |
| | | | 2E9W-A3 | LBD | Ig-like (1) |
| | | | 2E9W-A4 | - | Ig-like (1) |
| | | | 2E9W-A5 | - | Ig-like (1) |
| | FMS | P09581 | 3EJJ-B1 | - | Ig-like (2) |
| | | | 3EJJ-B2 | LBD | Ig-like (2) |
| | | | 3EJJ-B3 | LBD | Ig-like (2) |
| | | | FMS-B4 (AF2) | - | Ig-like (2) |
| | | | FMS-B5 (AF2) | - | Ig-like (2) |
| | PDGFRa | P16234 | 7LBF-C1 | LBD | Ig-like (3) |
| | | | 7LBF-C2 | LBD | Ig-like (3) |
| | | | 7LBF-C3 | LBD | Ig-like (3) |
| | | | PDGFRa-C4 (AF2) | - | Ig-like (3) |
| | | | PDGFRa-C5 25(AF2) | - | Ig-like (3) |
| | PDGFRb | P09619 | 3MJG-D1 | - | Ig-like (4) |
| | | | 3MJG-D2 | LBD | Ig-like (4) |
| | | | 3MJG-D3 | LBD | Ig-like (4) |
| | | | PDGFRb-D4 (AF2) | - | Ig-like (4) |
| | | | PDGFRb-D5 (AF2) | - | Ig-like (4) |
| | Flt3 | P36888 | 3QS9-E1 | LBD | Ig-like (5) |
| | | | 3QS9-E2 | LBD | Ig-like (5) |
| | | | 3QS9-E3 | LBD | Ig-like (5) |
| | | | 3QS9-E4 | - | Ig-like (5) |
| | | | 3QS9-E5 | - | Ig-like (5) |
| Axl | Axl | P30530 | 2C5D-A1 | LBD | Ig-like (6) |
| | | | 2C5D-A2 | LBD | Ig-like (6) |
| | | | Axl-A3 (AF2) | - | FN3 (6) |
| | | | Axl-A4 (AF2) | - | FN3 (6) |
| | Tyro3 | Q06418 | 1RHF-B1 | LBD | Ig-like (6) |
| | | | 1RHF-B2 | LBD | Ig-like (6) |

| | | | | | |
|---|---|---|---|---|---|
| | | | Tyro3-B3 (AF2) | - | FN3 [6] |
| | | | Tyro3-B4 (AF2) | - | FN3 [6] |
| | Mer | Q12866 | MER-C1 (AF2) | LBD | Ig-like [6, 7] |
| | | | MER-C2 (AF2) | LBD | Ig-like [6, 7] |
| | | | MER-C3 (AF2) | - | FN3 [6, 7] |
| | | | MER-C4 (AF2) | - | FN3 [6, 7] |
| Trk | TrkA | P04629 | 2IFG–A1 | - | Leucin-rich [8] |
| | | | 2IFG–A2 | - | Ig-like [8] |
| | | | 2IFG–A3 | LBD | Ig-like [8] |
| | TrkB | Q16620 | TrkB-B1 (AF2) | - | Leucin-rich [9] |
| | | | TrkB-B2 (AF2) | - | Ig-like [9] |
| | | | TrkB-B3 (AF2) | LBD | Ig-like [9] |
| | TrkC | Q16288 | TrkC-C1(AF2) | - | Leucin-rich [9] |
| | | | TrkC-C2 (AF2) | - | Ig-like [9] |
| | | | TrkC-C3 (AF2) | LBD | Ig-like [9] |
| Tie | Tie1 | P35590 | Tie1-A1 (AF2) | - | Ig-like [10] |
| | | | Tie1-A2 (AF2) | LBD | Ig-like [10] |
| | | | Tie1-A3 (AF2) | - | EGF [10] |
| | | | Tie1-A4 (AF2) | - | EGF [10] |
| | | | Tie1-A5 (AF2) | - | EGF [10] |
| | | | Tie1-A6 (AF2) | - | Ig-like [10] |
| | | | Tie1-A7 (AF2) | - | FN3 [10] |
| | | | Tie1-A8 (AF2) | - | FN3 [10] |
| | | | Tie1-A9 (AF2) | - | FN3 [10] |
| | TIE2 | Q15389 | 4K0V-B1 | - | Ig-like [11, 12] |
| | | | 4K0V-B2 | LBD | Ig-like [11, 12] |
| | | | 4K0V-B3 | - | EGF [11, 12] |
| | | | 4K0V-B4 | - | EGF [11, 12] |
| | | | 4K0V-B5 | - | EGF [11, 12] |
| | | | 4K0V-B6 | - | Ig-like [11, 12] |
| | | | 4K0V-B7 | - | FN3 [11, 12] |
| | | Q02763 | 5MYA-B8 | - | FN3 [11, 12] |
| | | | 5MYA-B9 | - | FN3 [11, 12] |

| | | | | | |
|---|---|---|---|---|---|
| ROR | ROR1 | Q01973 | ROR1-A1 (AF2) | Unk | Ig-like [13] |
| | | | ROR1-A2 (AF2) | Unk | Frizzled [13] |
| | | | ROR1-A3 (AF2) | Unk | Kringle [13] |
| | ROR2 | Q01974 | ROR2-B1 (AF2) | Unk | Ig-like [13] |
| | | | ROR2-B2 (AF2) | Unk | Frizzled [13] |
| | | | ROR2-B3 (AF2) | Unk | Kringle [13] |
| InsR | InsR | P06213 | 6PXV-A1 | LBD | Leucin-rich [14] |
| | | | 6PXV-A2 | LBD | Cysteine-rich [14] |
| | | | 6PXV-A3 | - | Leucin-rich [14] |
| | | | 6PXV-A4 | LBD | FN3 [14] |
| | | | 6PXV-A5 | - | FN3 [14] |
| | | | 6PXV-A6 | - | FN3 [14] |
| | IGF1R | P08069 | 6JK8-B1 | LBD | Leucin-rich [15] |
| | | | 6JK8-B2 | LBD | Cysteine-rich [15] |
| | | | 6JK8-B3 | - | Leucin-rich [15] |
| | | | 6JK8-B4 | LBD | FN3 [15] |
| | | | 6JK8-B5 | - | FN3 [15] |
| | | | 6JK8-B6 | - | FN3 [15] |
| | InsRR | P14616 | 7TYM-C1 | LBD | Leucin-rich [16] |
| | | | 7TYM-C2 | LBD | Cysteine-rich [16] |
| | | | 7TYM-C3 | - | Leucin-rich [16] |
| | | | 7TYM-C4 | LBD | FN3 [16] |
| | | | 7TYM-C5 | - | FN3 [16] |
| | | | 7TYM-C6 | - | FN3 [16] |
| PTK7 | PTK7 | Q13308 | PTK7-A1 (AF2) | Unk | Ig-like [17] |
| | | | PTK7-A2 (AF2) | Unk | Ig-like [17] |
| | | | PTK7-A3 (AF2) | Unk | Ig-like [17] |
| | | | PTK7-A4 (AF2) | Unk | Ig-like [17] |
| | | | PTK7-A5 (AF2) | Unk | Ig-like [17] |
| | | | PTK7-A6 (AF2) | Unk | Ig-like [17] |
| | | | PTK7-A7 (AF2) | Unk | Ig-like [17] |
| MET | MET | P08581 | 7MO7-A1 | LBD | Sema [18] |
| | | | MET-A2 (AF2) | - | Psl [18] |

| | | | | | |
|---|---|---|---|---|---|
| | | | MET-A3 (AF2) | - | Ig-like [18] |
| | | | MET-A4 (AF2) | - | Ig-like [18] |
| | | | MET-A5 (AF2) | - | Ig-like [18] |
| | RON | Q04912 | RON-B1 (AF2) | LBD | Sema [18] |
| | | | RON-B2 (AF2) | - | Psl [18] |
| | | | RON-B3 (AF2) | - | Ig-like [18] |
| | | | RON-B4 (AF2) | - | Ig-like [18] |
| | | | RON-B5 (AF2) | - | Ig-like [18] |
| RET | RET | P07949 | 6Q2O-A1 | - | Cadherin-like [19] |
| | | | 6Q2O-A2 | - | Cadherin-like [19] |
| | | | 6Q2O-A3 | - | Cadherin-like [19] |
| | | | 6Q2O-A4 | - | Cadherin-like [19] |
| | | | 6Q2O-A5 | LBD | Cysteine-rich [19] |
| ALK | ALK | Q9UM73 | ALK-A1(AF2) | - | Heparin binding domain (23 aa) [20] |
| | | | ALK-A2(AF2) | - | MAM domain 1 [20] |
| | | | ALK-A3(AF2) | - | LDL domain [20] |
| | | | ALK-A4(AF2) | - | MAM domain 2 [20] |
| | | | ALK-A6(AF2) | - | EGF [20] |
| | | | 7NX4-A5 | LBD | TNFL-GR (TG) ) [20] |
| | LTK | P29376 | 7NX0-C1 | LBD | TNFL-GR (TG) [20] |
| | | | LTK-C2 (AF2) | - | EGF [20] |
| Ryk | Ryk | P34925 | Ryk-A1 (AF2) | Unk | WIF [21] |
| ROS | ROS1 | P08922 | ROS-A2 (AF2) | Unk | FN3 [22] |
| | | | ROS-A3(AF2) | Unk | FN3 [22] |
| | | | ROS-A4 (AF2) | Unk | YWTD [22] |
| | | | ROS-A5(AF2) | Unk | FN3 [22] |
| | | | ROS-A6 (AF2) | Unk | YWTD [22] |
| | | | ROS-A7 (AF2) | Unk | FN3 [22] |
| | | | ROS-A8 (AF2) | Unk | FN3 [22] |
| | | | ROS-A9 (AF2) | Unk | YWTD [22] |
| | | | ROS-A10 (AF2) | Unk | FN3 [22] |
| | | | ROS-A11 (AF2) | Unk | FN3 [22] |

| | | | | | |
|---|---|---|---|---|---|
| | | | ROS-A12(AF2) | Unk | FN3 [22] |
| | | | ROS-A13 (AF2) | Unk | FN3 [22] |
| FGFR | FGFR1 | P11362 | 1CVS-A2 | LBD | Ig-like [23] |
| | | | 1CVS-A3 | LBD | Ig-like [23] |
| | | | FGFR1-A1 (AF2) | - | Ig-like [23] |
| | FGFR2c | P21802 | 2FDB-B2 | LBD | Ig-like [23] |
| | | | 2FDB-B3 | LBD | Ig-like [23] |
| | | | FGFR2-B1 (AF2) | - | Ig-like [23] |
| | FGFR3 | Q8NI16 | 3GRW-C2 | LBD | Ig-like [24] |
| | | | 3GRW-C3 | LBD | Ig-like [24] |
| | | | FGFR3-C1 (AF2) | - | Ig-like [24] |
| | FGFR4 | P22455 | 7YSW-D2 | LBD | Ig-like [25] |
| | | | 7YSW-D3 | LBD | Ig-like [25] |
| | | | FGFR4-D4(AF2) | - | Ig-like [25] |
| VEGFR | VEGFR1 | P17948 | 5T89-A1 | - | Ig-like [26] |
| | | | 5T89-A2 | LBD | Ig-like [26] |
| | | | 5T89-A3 | LBD | Ig-like [26] |
| | | | 5T89-A4 | - | Ig-like [26] |
| | | | 5T89-A5 | - | Ig-like [26] |
| | | | 5T89-A6 | - | Ig-like [26] |
| | | | VEGFR1-A7 (AF2) | - | Ig-like [26] |
| | VEGFR2 | P35968 | VEGFR2-B1 (AF2) | - | Ig-like [27] |
| | | | 2X1X-B2 | LBD | Ig-like [27] |
| | | | 2X1X-B3 | LBD | Ig-like [27] |
| | | | VEGFR2-B4 (AF2) | - | Ig-like [27] |
| | | | VEGFR2-B5 (AF2) | - | Ig-like [27] |
| | | | VEGFR2-B6 (AF2) | - | Ig-like [27] |
| | | | 3KVQ-B7 | - | Ig-like [27, 28] |
| | VEGFR3 | P35916 | 4BSK-C1 | - | Ig-like [29] |
| | | | 4BSK-C2 | LBD | Ig-like [29] |
| | | | VEGFR3-C3 (AF2) | LBD | Ig-like [29] |
| | | | VEGFR3-C4 (AF2) | - | Ig-like [29] |
| | | | VEGFR3-C5 (AF2) | - | Ig-like [29] |

| | | | | | |
|---|---|---|---|---|---|
| | | | VEGFR3-C6 (AF2) | - | Ig-like [29] |
| | | | VEGFR3-C7 (AF2) | - | Ig-like [29] |
| EphR | EphA1 | P21709 | EphA1-A1 (AF2) | LBD | Jelly-roll [30] |
| | | | EphA1-A2 (AF2) | - | Cysteine-rich [30] |
| | | | EphA1-A3 (AF2) | - | FN3 [30] |
| | | | EphA1-A4 (AF2) | - | FN3 [30] |
| | EphA2 | P29317 | 3FL7-B1 | LBD | Jelly-roll [30, 31] |
| | | | 3FL7-B2 | - | Cysteine-rich [30, 31] |
| | | | 3FL7-B3 | - | FN3 [30, 31] |
| | | | 3FL7-B4 | - | FN3 [30, 31] |
| | EphA3 | P29320 | EphA3-C1 (AF2) | LBD | Jelly-roll [30] |
| | | | EphA3-C2 (AF2) | - | Cysteine-rich [30] |
| | | | EphA3-C3 (AF2) | - | FN3 [30] |
| | | | EphA3-C4 (AF2) | - | FN3 [30] |
| | EphA4 | P54764 | 4M4R-D1 | LBD | Jelly-roll [30, 32] |
| | | | 4M4R-D2 | - | Cysteine-rich [30, 32] |
| | | | 4M4R-D3 | - | FN3 [30, 32] |
| | | | 4M4R-D4 | - | FN3 [30, 32] |
| | EphA5 | P54756 | EphA5-E1 (AF2) | LBD | Jelly-roll [30] |
| | | | EphA5-E2 (AF2) | - | Cysteine-rich [30] |
| | | | EphA5-E3 (AF2) | - | FN3 [30] |
| | | | EphA5-E4 (AF2) | - | FN3 [30] |
| | EphA6 | Q9UF33 | EphA6-F1 (AF2) | LBD | Jelly-roll [30] |
| | | | EphA6-F2 (AF2) | - | Cysteine-rich [30] |
| | | | EphA6-F3 (AF2) | - | FN3 [30] |
| | | | EphA6-F4 (AF2) | - | FN3 [30] |
| | EphA7 | Q15375 | EphA7-G1 (AF2) | LBD | Jelly-roll [30] |
| | | | EphA7-G2 (AF2) | - | Cysteine-rich [30] |
| | | | EphA7-G3 (AF2) | - | FN3 [30] |
| | | | EphA7-G4 (AF2) | - | FN3 [30] |
| | EphA8 | P29322 | EphA8-H1 (AF2) | LBD | Jelly-roll [30] |
| | | | EphA8-H2 (AF2) | - | Cysteine-rich [30] |
| | | | EphA8-HA3 (AF2) | - | FN3 [30] |

| | | | | | |
|---|---|---|---|---|---|
| | | | EphA8-H4 (AF2) | - | FN3 [30] |
| | EphA10 | Q5JZY3 | EphA10-I1 (AF2) | LBD | Jelly-roll [30] |
| | | | EphA10-I2 (AF2) | - | Cysteine-rich [30] |
| | | | EphA10-I3 (AF2) | - | FN3 [30] |
| | | | EphA10-I4 (AF2) | - | FN3 [30] |
| | EphB1 | P54762 | EphB1-J1 (AF2) | LBD | Jelly-roll [30] |
| | | | EphB1-J2 (AF2) | - | Cysteine-rich [30] |
| | | | EphB1-J3 (AF2) | - | FN3 [30] |
| | | | EphB1-J4 (AF2) | - | FN3 [30] |
| | EphB2 | P54763 | 7S7K-K1 | LBD | Jelly-roll [30, 33] |
| | | | 7S7K-K2 | - | Cysteine-rich [30, 33] |
| | | | 7S7K-K3 | - | FN3 [30, 33] |
| | | | 7S7K-K4 | - | FN3 [30, 33] |
| | EphB3 | P29320 | EphB3-L1 (AF2) | LBD | Jelly-roll [30] |
| | | | EphB3-L2 (AF2) | - | Cysteine-rich [30] |
| | | | EphB3-L3 (AF2) | - | FN3 [30] |
| | | | EphB3-L4 (AF2) | - | FN3 [30] |
| | EphB4 | P54760 | EphB4-M1 (AF2) | LBD | Jelly-roll [30] |
| | | | EphB4-M2 (AF2) | - | Cysteine-rich [30] |
| | | | EphB4-M3 (AF2) | - | FN3 [30] |
| | | | EphB4-M4 (AF2) | - | FN3 [30] |
| | EphB6 | O15197 | 7K7J-N1 | LBD | Jelly-roll [30, 34] |
| | | | 7K7J-N2 | - | Cysteine-rich [30, 34] |
| | | | 7K7J-N3 | - | FN3 [30, 34] |
| | | | EphB6-N4 (AF2) | - | FN3 [30, 34] |
| ErbB | EGFR | P00533 | 1IVO-A1 | LBD | Leucin-rich [35] |
| | | | 1IVO-A2 | - | EGF [35] |
| | | | 1IVO-A3 | LBD | Leucin-rich [35] |
| | | | EGFR-A4 (AF2) | - | EGF [35] |
| | HER2/ERbB2 | P04626 | 1N8Z-B1 | LBD | Leucin-rich [36] |
| | | | 1N8Z-B2 | - | EGF [36] |
| | | | 1N8Z-B3 | LBD | Leucin-rich [36] |
| | | | 1N8Z-B4 | - | EGF [36] |

| | | | | | |
|---|---|---|---|---|---|
| | HER3 | P21860 | 4LEO-D1 | LBD | Leucin-rich [37] |
| | | | 4LEO-D2 | - | EGF [37] |
| | | | 4LEO-D3 | LBD | Leucin-rich [37] |
| | | | 4LEO-D4 | - | EGF [37] |
| | HER4 | Q15303 | 2AHX-C1 | LBD | Leucin-rich [38] |
| | | | 2AHX-C2 | - | EGF [38] |
| | | | 2AHX-C3 | LBD | Leucin-rich [38] |
| | | | 2AHX-C4 | - | EGF [38] |
| DDR | DDR1 | Q08345 | 4AG4-A1 | LBD | Discoidin [39] |
| | | | 4AG4-A2 | LBD | Discoidin [39] |
| | DDR2 | Q16832 | 2WUH-B1 | LBD | Discoidin [40] |
| | | | DDR2-B2 (AF2) | LBD | Discoidin [39, 40] |
| MuSK | MusK | Q62838 | 2IEP-A1 | Unk | Ig-like [41] |
| | | | 2IEP-A2 | Unk | Ig-like [41] |
| | | | MusK-A3 (AF2) | Unk | Ig-like [42] |
| | | | 3HKL-A4 | Unk | Frizzled [42] |

**Supplementary Table 1. Structural domains in the 18 RTK subfamilies classified by functional and by structural features.** 245 individual structural domians, in each subfamiliy across the RTK superfamily. Domain identifiers are composed of PDB IDs and receptor names. Domains that bind ligands are marked as LBD, domains from orphan receptors are marked as Unk. Ig-like, Immunoglobulin-like domain, FN3, Fibronectin type III domain. The source for 3D fold classifications are in superscript.
